# The molecular framework of heterophylly in *Callitriche palustris* L. differs from that in other amphibious plants

**DOI:** 10.1101/2020.12.19.423571

**Authors:** Hiroyuki Koga, Mikiko Kojima, Yumiko Takebayashi, Hitoshi Sakakibara, Hirokazu Tsukaya

## Abstract

Heterophylly refers to the development of different leaf forms in a single plant depending on the environmental conditions. It is often observed in amphibious aquatic plants that can grow under aerial and submerged conditions. Although heterophylly is well recognized in aquatic plants, the associated developmental mechanisms and the molecular basis remain unclear. In this study, we analyzed heterophyllous leaf formation in an aquatic plant, *Callitriche palustris*, to clarify the underlying developmental and molecular mechanisms. Morphological analyses revealed extensive cell elongation and the rearrangement of cortical microtubules in the elongated submerged leaves of *C. palustris*. Our observations also suggested that gibberellin, ethylene, and abscisic acid regulate the formation of submerged leaves. However, the perturbation of one or more of the hormones was insufficient to induce the formation of submerged leaves under aerial conditions. Finally, we analyzed gene expression changes during aerial and submerged leaf development and narrowed down the candidate genes controlling heterophylly via transcriptomic comparisons, including a comparison with a closely related terrestrial species. We revealed that the molecular mechanism regulating heterophylly in *C*. *palustris* is associated with hormonal changes and diverse transcription factor gene expression profiles, suggesting differences from the corresponding mechanisms in previously investigated amphibious plants.

## Introduction

Plants have a remarkable variety of leaf forms (Tsukaya, 2018). How such diverse leaf forms evolved is a central question related to the evolution of land plants. Although leaf forms generally vary between species or sometimes between populations, in some species, individual plants can produce completely different leaf forms depending on the environmental conditions the shoot is exposed to (Arber, 1920; Allsopp, 1967; Sculthorpe, 1967). The ability to form different types of leaves depending on the environment is called heterophylly. Because an individual heterophyllous plant has distinct leaf developmental activities, it is useful for studying various modes of leaf development in a single genetic background.

Amphibious plants that can grow under aerial and submerged conditions are often used to investigate heterophylly, which is quite extensive in these species (Arber, 1920; Sculthorpe, 1967). Heterophylly is thought to be an important adaptive feature of amphibious plants, which usually grow under conditions in which water levels fluctuate seasonally or unexpectedly (Bradshaw, 1965). Amphibious plants often bear different types of leaves when their shoot apex is underwater to either rapidly escape from the submerged state or to adapt to underwater conditions. In the latter case, the leaves produced under submerged conditions are generally thin, filamentous, and sometimes highly branched, with a long and narrow leaf blade and/or a higher surface area relative to volume (Wells and Pigliucci, 2000). These characteristics are thought to be advantageous for surviving underwater. Because aquatic plants emerged from various lineages of land plants (Cook, 1999; Du and Wang, 2014), the subsequent heterophylly in response to submergence likely evolved independently in each lineage.

Previous studies revealed that phytohormones are involved in controlling heterophylly in aquatic plants (Wanke, 2011; Nakayama et al., 2017; Li et al., 2019). In many aquatic plants, gibberellic acid (GA) and ethylene promote the formation of submerged leaves, whereas abscisic acid (ABA) stimulates the formation of aerial leaves. Although the morphological basis of heterophylly has been well studied in various aquatic plants, the underlying molecular mechanisms remained unknown until recently. Specifically, Nakayama et al. (2014) clarified the molecular development related to heterophylly in the aquatic plant *Rorippa aquatica* (Brassicaceae), which forms branched dissected leaves when submerged. They revealed that GA negatively regulates dissected leaf development, which is correlated with both submergence and low temperatures, possibly by altering the expression patterns of class I KNOTTED1-LIKE HOMEOBOX (KNOX) genes in leaf primordia (Nakayama et al., 2014). In another study, in which *Ranunculus* species (Ranunculaceae) were used, heterophyllous leaf development was determined to be regulated by ethylene and ABA signaling that modifies the expression of leaf polarity genes, including those encoding KANADI and HD-ZIP III transcription factors (TFs) (Kim et al., 2018). These studies suggest that the molecular basis of heterophylly in aquatic plants varies among species. Hence, studying diverse species is important for elucidating plant adaptations to aquatic environments.

In this study, we analyzed leaf formation in *Callitriche* plants (known as water starworts). Members of the genus *Callitriche* are small flowering plants with a broad global distribution. It is a member of the Plantaginaceae, to which the well-studied snapdragon (*Antirrhinum majus*) belongs. *Callitriche* species are typically amphibious, able to grow both on land and in water (e.g., in freshwater ponds, streams, and rivers) (Erbar and Leins, 2004). This genus also comprises some terrestrial species that usually grow in moist soil. Although the sister genus of *Callitriche* is an amphibious genus, *Hippuris*, it was recently proposed that *Callitriche* has a terrestrial ancestor because the basal clade of the genus was revealed a terrestrial lifestyle (Ito et al., 2017). The other terrestrial species in the genus were likely derived from an amphibious ancestor. Thus, the evolution toward aquatic habitats and reversal of the terrestrial habitat may have occurred in the *Calltriche* lineage (Philbrick and Les, 2000; Philbrick and Jansen, 1991). Accordingly, *Callitriche* plants are suitable materials for studying the evolutionary mechanisms mediating adaptations to aquatic environments.

The significant heterophylly of some *Callitriche* species has long been recognized (e.g., Schenck, 1887), and many classic developmental and physiological studies have been conducted using *Callitriche* species to understand the mechanisms of aquatic adaptation. Regarding leaf development, several descriptive works and physiological experiments have been performed. For example, Jones described the development of leaves under both aerial and submerged conditions using *C*. *intermedia* (= *C*. *hamulata*), *C*. *stagnalis*, and *C*. *obtusangula* (Jones, 1952, 1955). More recently, Deschamp and Cooke reported dimorphic leaf development in *C*. *heterophylla* at the cellular level and suggested the involvement of phytohormones and turgor pressure in the control of heterophylly in this plant (Deschamp and Cooke, 1984, 1983, 1985). Besides, we recently reported that *C. palustris*, a close relative of *C. heterophylla*, showed extensive heterophylly in response to submergence even in the laboratory (Koga et al., 2020). We documented the aerial and submerged leaf developmental processes in this species, and defined the stages of leaf development on the basis of advances in research regarding vascular and stomatal development and cell proliferation activities (Koga et al., 2020). We revealed that submerged leaf formation in *C. palustris* is characterized by differential cell proliferation (in terms of direction), extensive cell elongation, and decreases in the number of stomata and veins as well as in cuticle thickness. However, the mechanistic aspects of heterophylly, such as hormonal and genetic controls, were not analyzed. Here, we observed the cellular changes, hormonal effects, and changes in gene expression associated with heterophylly in this plant. We also report a potential regulatory gene set for dimorphic leaf formation in this species identified through comparative transcriptomic analyses among plants exposed to pharmacological treatments, as well as with the closely related terrestrial species *C. terrestris*.

## Results

### Differential cellular expansion in the heterophylly of *C. palustris*

A previous study reported that *C*. *palustris* exhibits extensive heterophylly in response to submergence (Schenck, 1887), and we showed that this heterophylly is reproducible under scalable experimental conditions (Koga et al., 2020). The plant forms ovate leaves when the shoots are exposed to aerial conditions, whereas it forms ribbon-like leaves when the shoots are submerged in water (Figure 1A, B). Additionally, the heterophylly of *C*. *palustris* is characterized by a decrease in the number of veins and the suppression of stomatal development in submerged leaves. The significantly narrower and longer submerged leaves were suggested to be the result of differential cellular arrangement and differential cell expansion (Koga et al., 2020). Because of the differential cell expansion, the aerial leaves were composed of jigsaw puzzle-like pavement cells and round palisade cells, whereas the cells of both the epidermal and mesophyll layers in submerged leaves were elongated along the proximal–distal axis (Figure 1C–F). A quantitative analysis of cell size and shape indicated that the pavement cells of submerged leaves had lower solidity values or were less lobed (Figure 1H; Supplemental Figure 1) and were significantly more elongated (Figure 1I) than the corresponding cells of aerial leaves. Similarly, the palisade cells of submerged leaves were less circular and much more elongated than the palisade cells of the aerial leaves (Figure 1K, L; Supplemental Figure 1). Despite the apparent differences in cell shape, the projected cell areas on the paradermal plane of both epidermal and palisade cells differed only slightly between the leaf forms (Figure 1G, J).

**Figure 1.**
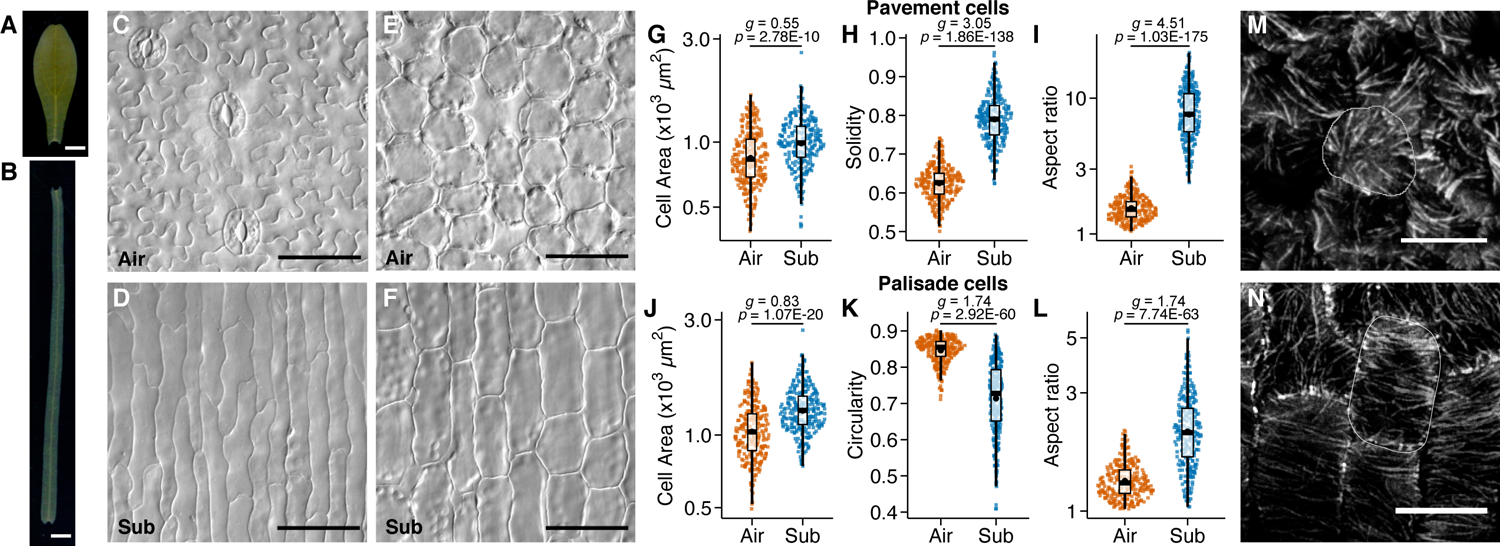
*Callitriche palustris* leaf cell morphology. (A, B) *C*. *palustris* leaves grown under aerial (A) and submerged conditions (B). (C) Epidermis of an aerial leaf. (D) Epidermis of a submerged leaf. (E) Sub-epidermal palisade layer of an aerial leaf. (F) Sub-epidermal palisade layer of a submerged leaf. (G–L) Plots of the indices for pavement cells (G–I, n = 270 each) and palisade cells (J–L, n = 270 each): cell area (G, J), solidity (H), circularity (K), and the aspect ratio of the best fit ellipse (I, L). Black points represent the means. *P*-values calculated by Welch’s *t*-test and Hedge’s *g* calculated to indicate the effect size are provided. Data were obtained for nine leaves from three biological replicates for each condition. (M, N) Immunofluorescence images of tubulin in the palisade cells of a young aerial leaf (M) and a young submerged leaf (N). A representative cell is outlined. Scale bars: (A, B) 1 mm, (C–F) 50 µm, and (M, N) 10 µm.

The directional expansion of plant cells is usually associated with the directional orientation of cell wall cellulose microfibers, which is controlled by the orientation of cortical microtubules (cMTs) (Preston, 1974; Green, 1980; Gunning and Hardham, 1982; Lloyd, 1991). We observed that the cMTs of the palisade cells in young (approximately 1 mm long) submerged leaves were oriented perpendicular to the axis of cell elongation, whereas they were not well organized in the palisade cells of young aerial leaves (Figure 1M, N). Therefore, the differences in cell shapes are likely due to the regulation of cMTs and the subsequent cellulose fiber orientation, resulting in directional cell expansion in submerged leaves.

### Effects of hormones on the formation of submerged leaves

As summarized in the Introduction, previous studies in diverse aquatic plants have revealed that phytohormones are generally involved in controlling heterophylly in response to submergence. In *C*. *heterophylla*, dimorphic leaf development is reportedly related to GA and ABA contents (Deschamp and Cooke, 1983). Ethylene is also a key regulator of the submergence response in various aquatic plants (Cox et al., 2004; Jackson, 2008). To confirm the effects of these phytohormones in *C*. *palustris*, we conducted hormone perturbation experiments.

After determining the effective concentrations of inhibitors or hormones (Supplemental Figure 2), we examined the phenotypes of mature leaves from submerged plants treated with these chemicals at the minimal effective concentrations (Figure 2, Supplemental Figure 4). When the plants were grown in water containing AgNO_3_, which inhibits ethylene signaling, they formed aerial-like leaves of a slightly shorter length (Figure 2B, H). After adding uniconazole P, an inhibitor of GA synthesis, to the water, the plants also produced leaves that were shorter and broader than typical submerged leaves (Figure 2C, H). A similar tendency was also observed when a different GA synthesis inhibitor, paclobutrazol, was added to the water (Supplemental figure 2). The inhibition of submerged-type leaf development by uniconazole P was recovered by adding GA_3_ (Figure 2G, H), implying GA production is required for the elongated leaf form. Additionally, plants grown in water containing GA_3_ exhibited an enhanced submerged leaf phenotype, with much longer and narrower leaves than normal, regardless of whether plants were also treated with uniconazole P (Figure 2E, G, H). On the other hand, combined treatment with AgNO_3_ and GA_3_ did not induce complete submerged-type leaves, although the leaves were longer and narrower than aerial leaves (Figure 2 F, H). Furthermore, the formation of submerged leaves was also inhibited when plants were grown in water containing ABA (Figure 2D, H). Both AgNO_3_ and ABA treatment drastically increased stomata density to the extent of normal aerial leaves, whereas uniconazole P and the combination of AgNO_3_ and GA_3_ led to slight increases. In the other GA_3_ treatment conditions, stomatal densities remained at the level of normal submerged leaves (Figure 2J).

**Figure 2.**
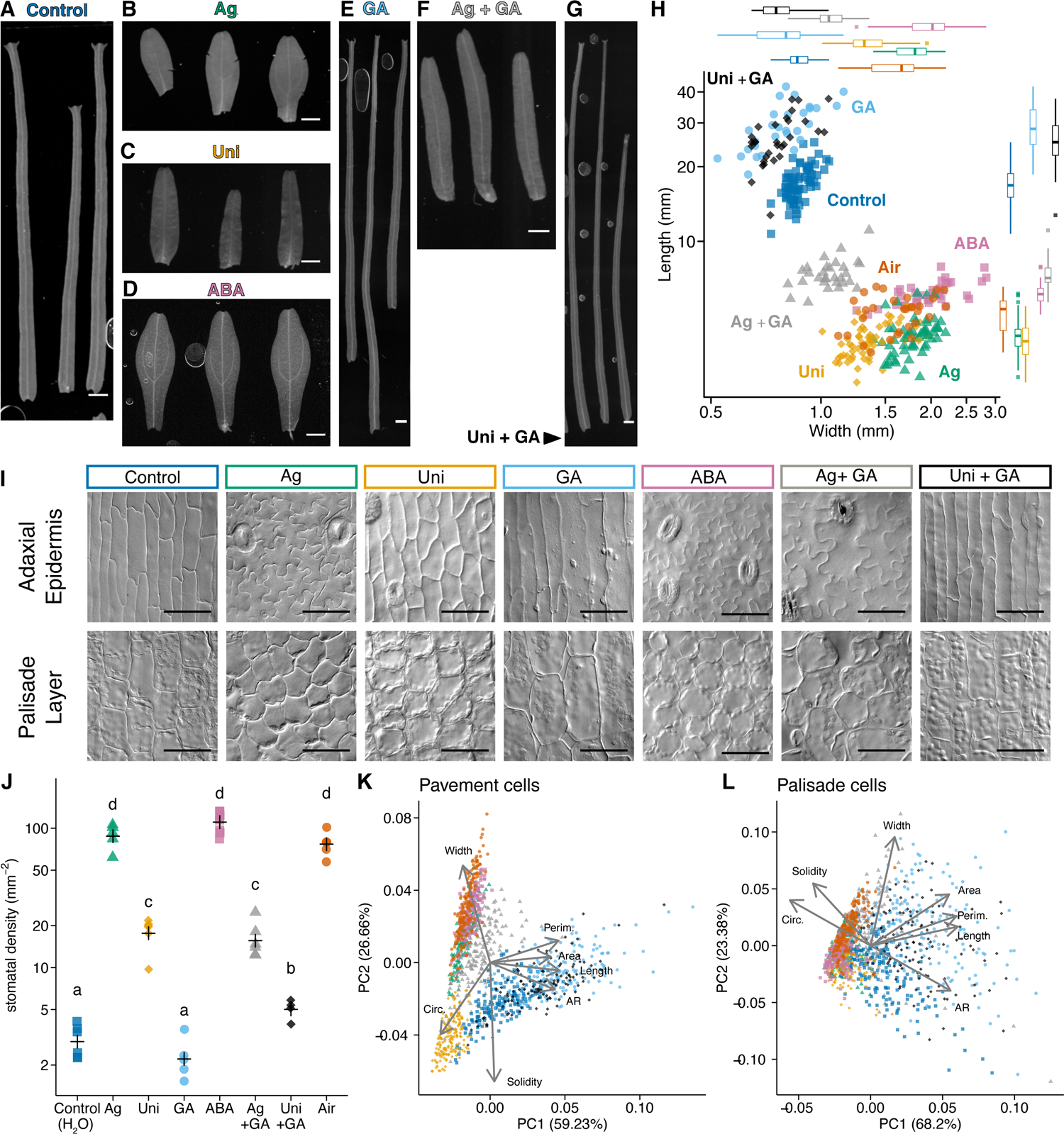
Effects of phytohormone inhibitors and phytohormones on *C. palustris* leaf development under submerged conditions. (A–G) Leaf forms of the plants treated with phytohormone inhibitors and/or phytohormones under submerged conditions: (A) control, (B) 10^−6^ M AgNO_3_, (C)10^−7^ M uniconazole P, (D) 10^−7^ M ABA, (E) 10^−6^ M GA_3_, (F) 10^−6^ M AgNO_3_ + 10^−6^ M GA_3_, and (G) 10^−7^ M uniconazole P + 10^−6^ M GA_3_. (H) Length–width plot of mature leaves. Each point represents one leaf. A total of 70 (for control) or 26–40 (for the others) mature leaves collected from 3-5 biological replicates were measured for each treatment. (I) Images of the adaxial epidermis (top) and sub-epidermal palisade layer (bottom) of the mature leaves. (J) Plots of stomatal density. Crosses represent the means. Different letters denote significant differences among treatments (Tukey’s test, *P* < 0.05; n = 4-6). (K, L) Principal component analysis plots of adaxial pavement cells (K) and palisade cells (L). Each point represents one cell (n = 120 or 180 for each treatment). Data were collected for four or six leaves from three biological replicates, except for the ABA treatment (six leaves from two replicates). Loading plots are overlapped on the graphs. Data for the leaves or the cells of the aerial control in Figure 3 (denoted as Air) are included in every plot for comparison purposes. Scale bars: 50 µm.

Regarding cell shape analyses, we observed and measured several cellular indices (i.e., cell area, perimeter, circularity, aspect ratio of the best fit ellipse [AR], solidity, width, and length; Supplemental Figure 1) to discriminate shapes among states (Figure 2K, L, Supplemental Figures 5, 6). In the leaves treated with AgNO_3_, ABA, and AgNO_3_ + GA_3_, the epidermal pavement cells and palisade cells were similar in shape to those of aerial leaves, although the cells in the AgNO_3_ + GA_3_-treated leaves expanded much more than the cells of leaves treated with AgNO_3_ alone (Figure 2I, K, L). These cells had a lower aspect ratio and shorter length than the cells of plants grown under control conditions (Supplemental figure 5), presumably representing their complex, jigsaw puzzle-like cellular shape. In uniconazole P-treated leaves, highly circular and relatively simple pavement cells were detected, indicating that the cells were less complex than those of aerial leaves, but they were not elongated, unlike the corresponding cells of submerged leaves (Figure 2I, K, L). Following the GA_3_ and uniconazole P + GA_3_ treatments, the leaf cells were highly expanded, and their shapes were similar to those of control leaf cells (Figure 2I, K, L). These observations suggest that inhibiting GA signaling alone is insufficient to block submerged-type cell differentiation, resulting in an intermediate leaf phenotype.

### Effects of hormones on the formation of aerial leaves

We next evaluated the effects of hormone applications on leaves under aerial conditions to determine whether GA or ethylene signals alone or combined are sufficient to trigger the formation of submerged leaves (Figure 3, Supplemental Figures 4, 5). Because an inhibitor of ethylene signaling prevented submerged leaves from forming, we grew the plants in medium containing ACC, which is an ethylene precursor, thereby enabling the activation of ethylene signaling even under aerial conditions. In this case, the plants exhibited a dwarf phenotype, which suggests strong inhibition of plant growth by ethylene signaling (Supplemental Figure 3). This phenotype was seemingly the opposite of that of submerged plants. Meanwhile, the plants produced slightly narrower leaves than controls, but it was insufficient for the development of submerged leaves under aerial conditions (Figure 3B, E, Supplemental Figure 4). To verify these results, we exposed plants to ethylene gas, which produced the same results as those obtained for the plants grown in ACC-containing medium (Supplemental Figure 3).

**Figure 3.**
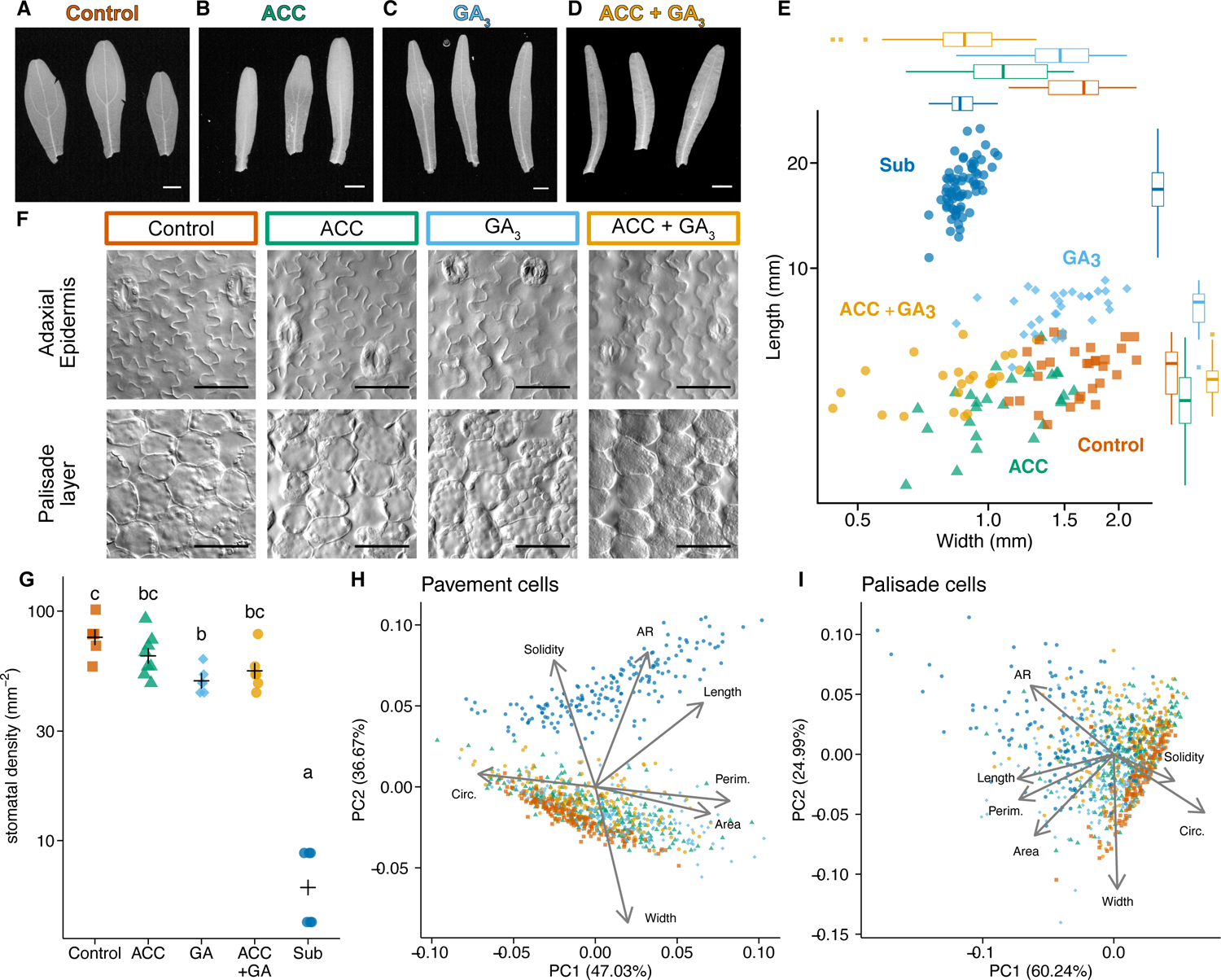
Effects of ACC and GA_3_ on *C*. *palustris* leaf development under aerial conditions. (A–D) Leaf forms of the plants treated with phytohormones under aerial conditions: (A) control, (B) 10^−5^ M ACC, (C) 10^−5^ M GA_3_, and (D) 10^−5^ M ACC + 10^−5^ M GA_3_. (E) Length–width plot of mature leaves. Each point represents one leaf. 25-30 mature leaves collected from 3 biological replicates were measured for each treatment. Points denoted by “Sub” represent the untreated submerged leaves shown in Figure 2 and are included for comparison purposes. (F) Images of the adaxial epidermis (top) and sub-epidermal palisade layer (bottom) of the mature leaves. (G) Plots of stomatal density. Crosses represent the means. Different letters denote significant differences among treatments (Tukey’s test, *P* < 0.05, n=4-8). (H, I) Principal component analysis plots of adaxial pavement cells (H) and palisade cells (I). Each point represents one cell (n = 180 for each treatment). Data were collected for six leaves from three biological replicates. Loading plots are overlapped on the graphs. Data for the leaves or the cells of the submerged control in Figure 2 (denoted as Sub) are included in every plot for comparison purposes. Scale bars: 50 µm.

**Figure 4.**
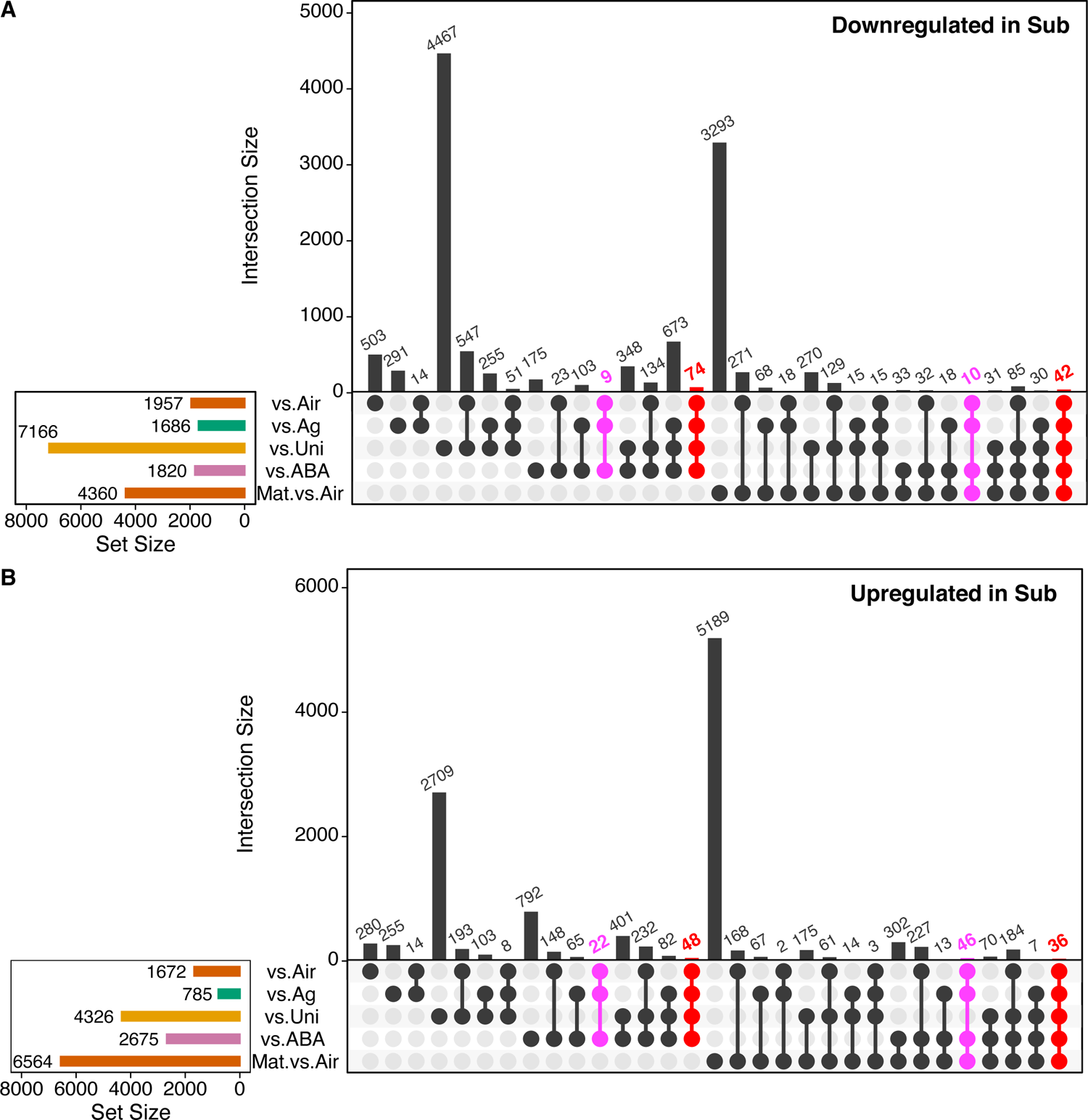
Comparative transcriptome analyses of *C. palustris*. UpSet plots present the overlapping DEGs in five comparisons. Gene sets that exhibited consistent changes among the comparisons of aerial, AgNO_3_-treated, and ABA-treated leaf primordia are colored (i.e., shared DEGs). (A) UpSet plot of DEGs downregulated in the submerged control. (B) UpSet plot of DEGs upregulated in the submerged control.

When GA_3_ was added to the medium, the plants elongated and developed leaves that were longer than those of the control plants, but GA_3_ did not induce the formation of normal submerged leaves (Figure 3C, E). Similar phenotypes were also induced by spraying plants with a GA_3_ solution (Supplemental Figure 3).

Finally, the plants were grown in medium containing both ACC and GA_3_ to activate ethylene and GA signaling under aerial conditions. However, this treatment resulted in slightly narrower leaves, but failed to induce the formation of submerged leaves (Figure 3D, E, Supplemental Figure 4). Stomatal densities decreased in response to GA_3_, which suggests that the GA signal negatively regulates stomatal formation (Figure 3G). However, the stomatal density was still higher than that of the submerged leaves. Substantial cellular changes in the aerial and submerged leaves were not observed following the application of hormones, although slight elongations and narrowing were detected that were consistent with the leaf forms (Figure 3F, H, I, Supplemental Figures 5, 6). Previous research on other aquatic plants proved that some submerged-like leaf phenotypes, such as a significantly narrow or compounded leaf form or extensive decreases in stomatal density, can be induced by a simple hormonal perturbation in many cases (Kuwabara et al., 2001; Sato et al., 2008; Nakayama et al., 2014; Kim et al., 2018; Horiguchi et al., 2019; Li et al., 2017). Therefore, our results suggest that the extent of the hormonal contribution to heterophylly differs between *C. palustris* and previously investigated amphibious plants.

**Figure 5.**
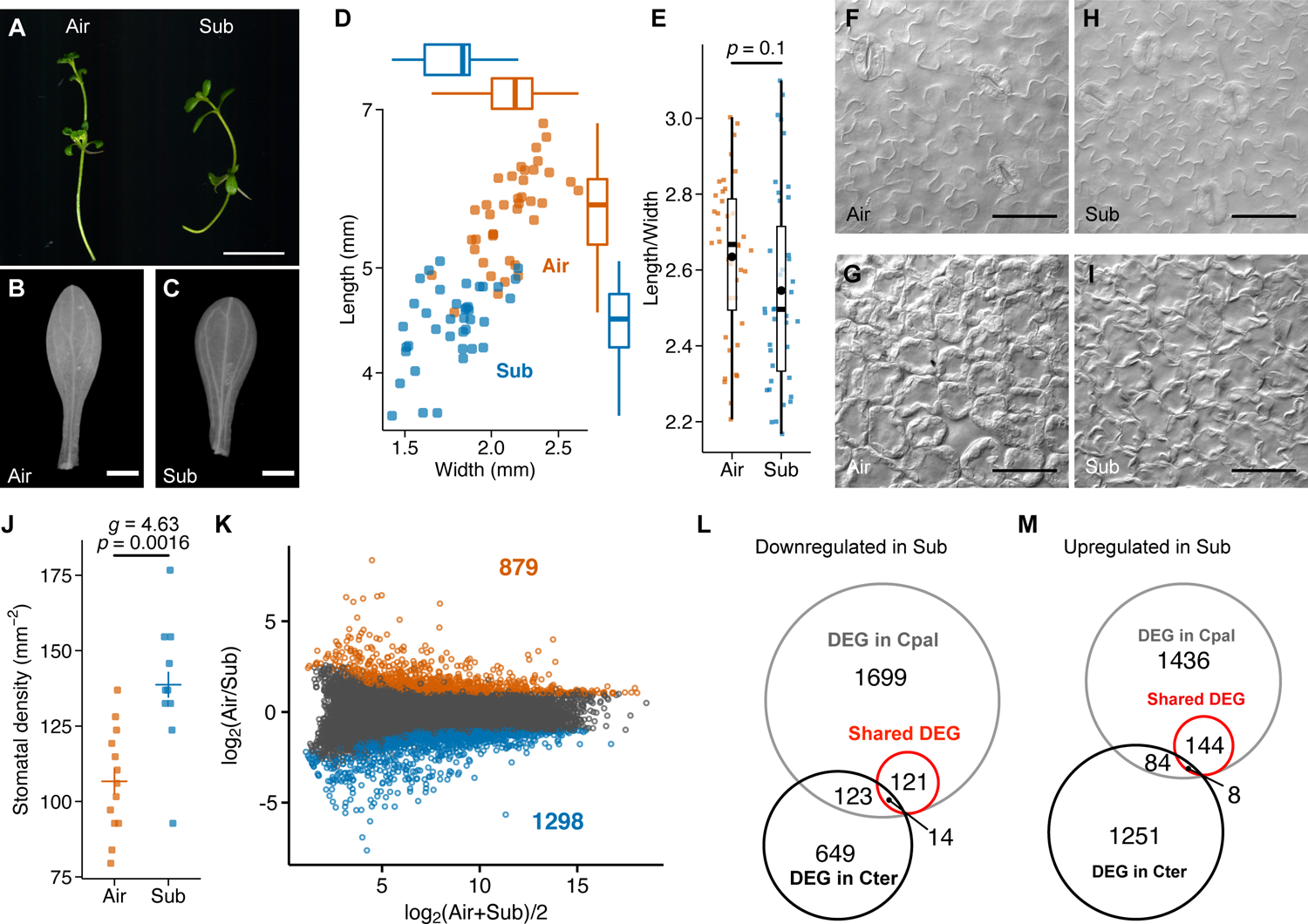
Submergence response of the terrestrial species *Callitriche terrestris*. (A) Images of shoots grown under aerial (left) and submerged (right) conditions. (B, C) Leaves from aerial (B) and submerged (C) shoots. (D) Length–width plots of mature leaves from both conditions. (E) Plots of length/width ratios for the mature leaves (n = 44 each, from 3 biological replicates). The *P* value was calculated by Welch’s *t*-test. (F–I) Images of the epidermal (E) and palisade layer (F) cells of a leaf from an aerial plant and the epidermal (G) and palisade layer (H) cells of a leaf from a submerged plant. (J) Plots of stomatal densities of mature leaves (n=13, 10, from 3 biological replicates). The *P* value calculated by Welch’s *t*-test and Hedge’s *g* calculated to indicate the effect size are provided. (K) MA plot for transcriptome analyses of the *C*. *terrestris* developing leaves. Red and blue points represent the downregulated and upregulated DEGs, respectively, under submerged conditions (FDR < 0.05 and log_2_(fold-change) > 1). The numbers of upregulated and downregulated DEGs are indicated. (L, M) Venn diagrams of *C. palustris* orthologs identified as DEGs in either *C. palustris* or *C. terrestris*.

**Figure 6.**
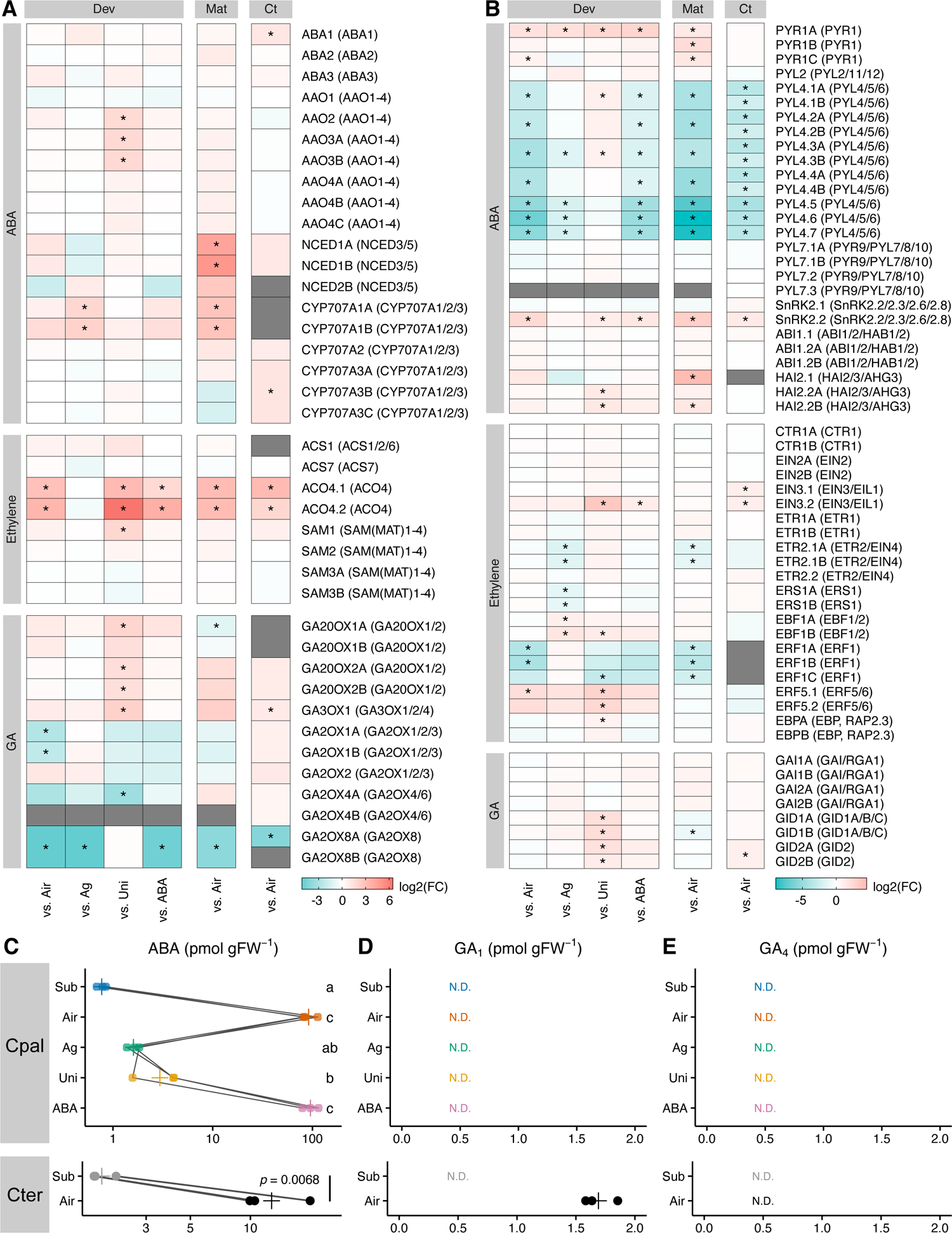
Hormone-related gene expression patterns and hormone contents in the shoot tips. (A, B) Expression-level changes to (A) putative hormone metabolism gene orthologs and (B) hormone signaling gene orthologs based on the RNA-seq data. The heatmaps present the mean log_2_(fold-change) values, relative to the corresponding levels in the submerged controls. Asterisks indicate significant differences (FDR < 0.05, expression-level log_2_(fold-change) > 1). Decreasing values reflect the upregulated expression in the submerged control. Gray panels indicate the gene, or its expression, was undetectable. Dev: comparisons among *C. palustris* leaf primordia, Mat: comparison between *C. palustris* mature leaves, and Ct: comparison between *C. terrestris* leaf primordia. The column on the right presents the names of *Callitriche* genes and the Arabidopsis orthologs determined using OrthoFinder (in parentheses). (C–E) Hormone contents: (C) ABA, (D) GA_1_, and (E) GA_4_ in the shoot tips of *C. palustris* (top) and *C. terrestris* (bottom). Different letters indicate significant differences among treatments (Tukey’s test, *P* < 0.05). For comparisons involving *C. terrestris*, the *P*-value was calculated by Welch’s *t*-test. N.D.: not detected.

### Gene expression patterns associated with different leaf developments

To address the molecular aspects of heterophylly in *C*. *palustris*, we performed a comprehensive mRNA sequencing analysis. Because the genome of the species has not been sequenced, we first reconstructed the transcriptome using RNA extracted from whole plants grown under aerial and submerged conditions (Supplemental Table 1). The data revealed many more genes in *C*. *palustris* than in known model species, possibly because the incomplete assembly resulted in the fragmentation of single genes and because of genome duplication events in *C*. *palustris*, which is supposedly a tetraploid species (n = 10, x = 5) (Philbrick, 1994; Prančl et al., 2014).

Regarding gene expression analyses, we extracted RNA from leaf primordia shorter than 500 µm, which were collected from three individuals grown under either aerial or submerged conditions. This growth stage was recently confirmed to precede or coincide with the initiation of cell differentiation under both conditions (Koga et al., 2020). We obtained single-read sequences (Supplemental Data Set 1) and found 4,275 differentially expressed genes (DEGs) between developing aerial and submerged leaves (false discovery rate [FDR] < 0.05 and expression-level log_2_(fold-change) > 1). We further analyzed gene expression patterns in the primordia submerged in water containing AgNO_3_, uniconazole P, or ABA, which inhibit the formation of submerged leaves. A total of 2,933, 5,508, and 16,517 genes were identified as DEGs in the AgNO_3_-, ABA-, and uniconazole P-treated leaf primordia, respectively, relative to the corresponding expression levels under normal submerged conditions (Figure 4). Because the leaves resulting from these treatments were significantly shorter and wider than the leaves formed under normal submerged conditions, we speculated that there is a key gene set that promotes the formation of submerged leaves among the common DEGs in the four comparisons. Interestingly, the whole transcriptome profiles of the treated primordia differed from that of the aerial leaf primordia despite their similar appearance (Supplemental Figure 7), indicating that the number of key gene set for differential development could be relatively a small fraction of the whole DEGs. Indeed, a total of 200 genes were identified as common DEGs among the four comparisons (Figure 4, red). The submerged plants treated with AgNO_3_ and ABA, but not uniconazole P, produced leaves that were almost the same as aerial leaves, implying the DEGs common to the AgNO_3_-treated, ABA-treated, and aerial leaf primordia, but not the uniconazole P-treated leaf primordia, include genes that are important for the development of submerged leaves. As such category, 87 genes were identified (Figure 4, magenta). Taken together, we narrowed down the important candidate genes for differential leaf development to 287 genes.

To determine whether the genes were differentially expressed only in developing leaves, we also compared the transcriptomes of mature aerial and submerged leaves. Of the 287 candidate genes, 153 were developmental stage-specific DEGs, whereas 134 were identified as DEGs in both developing and mature leaves (Figure 4). Although both types of DEGs may be important for heterophylly, those common to developing and mature leaves are likely related to general functions, including responses to environmental cues or signaling, whereas the DEGs specific to developing leaves are likely related to the regulation of cell division and differentiation.

### Analyses of non-heterophyllous *C*. *terrestris*

Although most *Callitriche* species are aquatic or amphibious, some are terrestrial, including *C*. *terrestris*, which grows in a relatively dry environment (Fassett, 1951). Phylogenetic analyses indicated that *C*. *terrestris* belongs to a sister clade of an amphibious clade that includes *C*. *palustris* and *C*. *heterophylla* (Philbrick and Les, 2000). Because the outer clade of this group also comprises amphibious species, *C. terrestris,* and other terrestrial species in this clade are believed to have an amphibious ancestor (Philbrick and Les, 2000). Thus, heterophylly in response to submergence likely weakened or was lost in these species during the transition to terrestrial habitats. To test this assumption, we collected *C*. *terrestris* plants and established a culturing system in our laboratory. The *C*. *terrestris* plants grew well in a growth chamber in either soil or Murashige and Skoog (MS) medium under sterile conditions. More specifically, in MS medium, the plants grew under both aerial and submerged conditions, but there were no major differences in leaf forms (Figure 5A–C). Compared with the aerial leaves, the submerged leaves were slightly shorter and narrower, which resulted in a decrease in the leaf area, but not in the leaf index (Figure 5D, E). Additionally, there were no differences in the leaf cell shapes (Figure 5F–I). Although stomatal density was rather increased in the submerged condition, considering the decrease of leaf area of the submerged plants, stomatal development was apparently not significantly affected by submergence. Accordingly, we concluded that *C*. *terrestris* does not exhibit heterophylly in response to submergence, unlike *C*. *palustris*.

Because submergence did not induce drastic changes in the *C*. *terrestris* leaf form, a comparison of gene expression patterns between *C. terrestris* and *C*. *palustris* was expected to provide relevant information regarding the key regulators of heterophylly. Therefore, we also analyzed the *C*. *terrestris* transcriptome. We first reconstructed the *C*. *terrestris* transcriptome using RNA-seq data for whole plants under both aerial and submerged conditions (Supplemental Table 1). To compare the gene expression patterns of *C. terrestris* with those of *C*. *palustris*, we analyzed orthologous relationships among putative coding genes. Because *C*. *terrestris* is a diploid species (n = 5; Philbrick 1994), we expected a one-to-many relationship with the orthologous genes in the tetraploid *C*. *palustris*. We identified 19,645 *C*. *terrestris* genes that were orthologous to 34,083 *C*. *palustris* genes (Supplemental Table 1). More precisely, 9,396 *C. terrestris* genes had a one-to-one relationship with *C. palustris* genes, whereas 10,249 genes were associated with 24,687 *C. palustris* genes having a one-to-multi relationship (Supplemental Figure 8). We subsequently sequenced the RNA extracted from the young leaves (< 500 µm long) collected from *C*. *terrestris* plants grown under aerial or submerged conditions. A total of 2,177 genes were differentially expressed between the developing leaves of *C*. *terrestris* plants grown under aerial and submerged conditions (Figure. 5K). Notably, the enriched gene ontology (GO) terms for the *C*. *palustris* and *C*. *terrestris* DEGs between the aerial and submerged leaf primordia were largely unshared (Supplemental Figure 9; Supplemental Data Set 2). Consistent with this result, we detected relatively few overlapping DEGs (Figure 5L, M).

### Expression-level changes in genes involved in hormone signaling

To examine the hormonal contribution to heterophylly from a gene expression aspect, we extracted the expression profiles of orthologous genes related to ethylene, GA, and ABA biosynthesis, inactivation, and signal transduction from the RNA-seq data (Figure 6A, B). Additionally, we quantified the contents of ABA, GAs, and other hormones in the shoots of *C. palustris* with hormone/inhibitors treatment as well as in the shoots of *C. terrestris* (Figure 6C-E, Supplemental Data Set 3). In *C. palustris*, the genes encoding proteins related to ABA biosynthesis (ABA1–3, AAOs, and NCEDs) tended to be highly expressed in mature aerial leaves, but not in primordia. The expression of genes encoding some ABA-inactivation enzymes (CYP70As) was also repressed in the *C. palustris* mature submerged leaves. Regarding *C. terrestris*, for which only the primordia were examined, there were no drastic changes in the expression of ABA biosynthesis genes like in the *C. palustris* primordia. Meanwhile, the ABA content decreased substantially in the submerged shoots (Figure 6C), consistent with the downregulation expression of NCED-encoding genes in the mature leaves. Similar results were obtained for the *C. terrestris* shoots. Among the signaling-related genes, there were notable changes in the expression of ABA receptor orthologs. Specifically, the expression of a *PYR1* ortholog was downregulated in the submerged leaves of *C. palustris*, whereas the expression of *PYL4–6* orthologs was upregulated in the submerged leaves of both species (Figure 6B).

Of the ethylene biosynthesis genes, the expression levels of the *ACO4* orthologs were clearly downregulated in the submerged control, relative to the corresponding levels in the aerial, uniconazole P-treated, and ABA-treated *C. palustris* and *C. terrestris* samples, suggesting ethylene biosynthesis was repressed. Considering this downregulated expression was not observed following the AgNO_3_ treatment, it might be the result of activation of ethylene signaling (i.e., negative feedback). Ethylene cannot diffuse effectively from plant tissues underwater. Hence, ethylene signaling was likely activated in the plants, even if the related biosynthesis genes were expressed at low levels. The upregulated expression of *ERF1* orthologs in the submerged primordia, but not in AgNO_3_-treated primordia, may reflect the activation of ethylene signaling (Solano et al., 1998).

Considering GA signaling is required for submerged leaf formation (Figure 2), it was surprising that we were unable to detect the activation of GA biosynthesis based on gene expression levels (Figure 6A). Instead, the expression of some orthologous genes related to GA deactivation (*GA2OX* genes) was upregulated under submerged conditions. Additionally, we did not detect active forms of GAs in *C. palustris* shoots under any condition, indicative of a lack of drastic changes in GA contents (Figure 6D, E). In contrast, in *C. terrestris*, the GA_1_ content decreased in submerged shoots (Figure 6D). Assuming that *GA2OX* expression levels are affected by the negative feedback regulation of GA signaling like in Arabidopsis (Thomas et al., 1999), GA signaling may be activated in submerged plants even at an extremely low GA level, possibly because of increased sensitivity. However, *GID1* receptor genes were similarly expressed under aerial and submerged conditions (Figure 6B).

### Gene expression patterns related to heterophyll**y**

We subsequently focused on the genes likely associated with heterophylly. Previous research on the molecular response to submergence proved that a subfamily of ERF TFs (i.e., group VII ERFs) contributes to the viability or the morphological changes of some plants under submerged condition (VEEN et al., 2014; Fukao et al., 2006; Xu et al., 2006; Licausi et al., 2011; Hattori et al., 2009). In Arabidopsis, the HRE1/2 and RAP2.12 TFs in this group function under low-oxygen stress conditions (i.e., hypoxia) to enhance flooding tolerance (Licausi et al., 2010). We detected three orthologous group VII ERF TF genes in the *Callitriche* transcriptomes. Of the three group VII ERF TF genes in *C*. *palustris*, two (*RAP2.2* and *RAP2.3*) were almost constitutively expressed under all conditions like *RAP2.12* of Arabidopsis, whereas the expression of the other orthologous gene (*HRE2*) was upregulated under submerged conditions, although not in the AgNO_3_-treated primordia (Figure 7A). The same tendency was observed for the orthologs in *C*. *terrestris* (Figure 7A).

**Figure 7.**
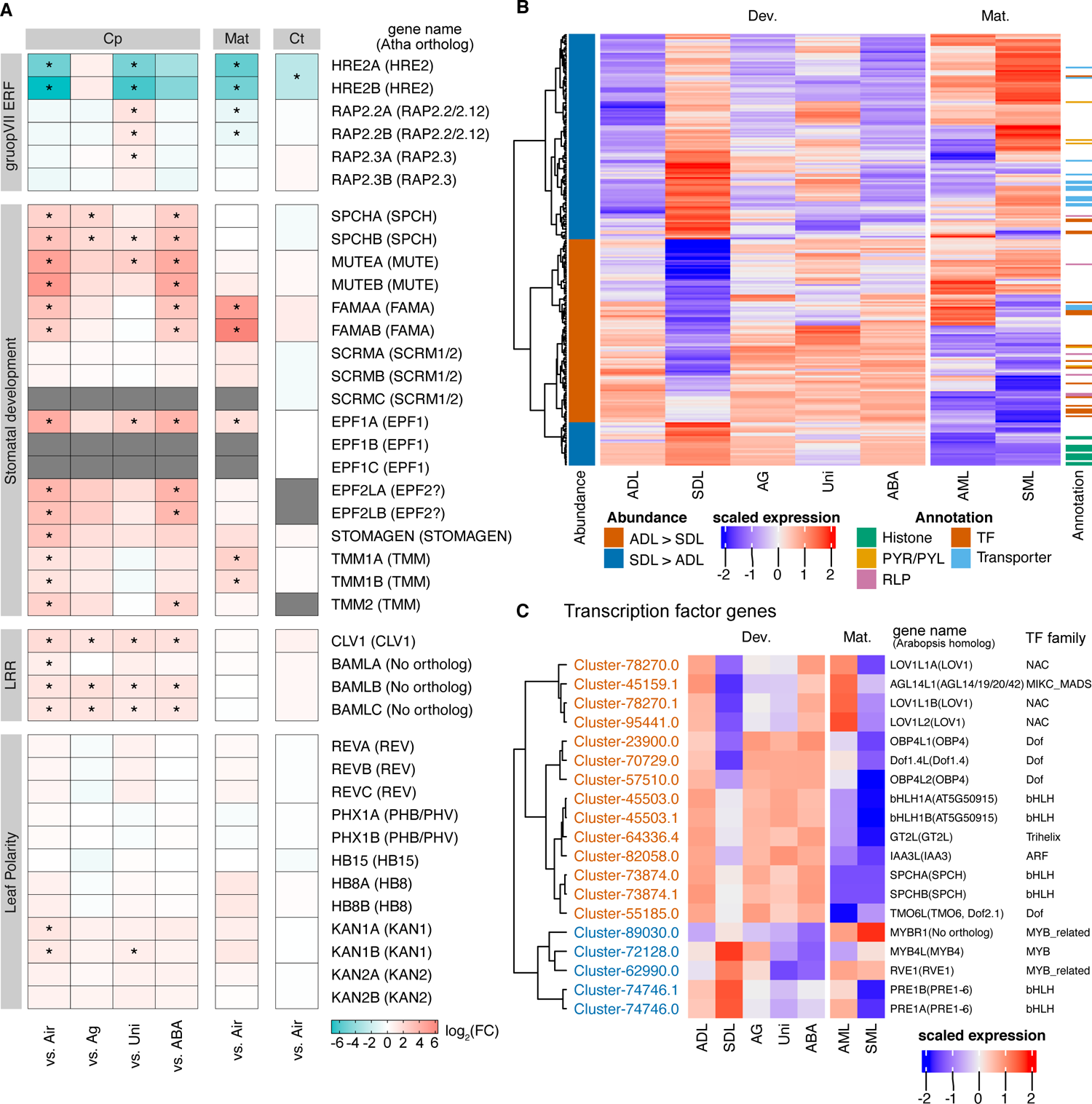
Expression patterns of the notable genes and the candidate genes. (A) Heatmaps of expression-level changes to some genes of interest based on the RNA-seq data. The data are presented in the same manner as in Figure 6A, B. (B) Candidate gene expression patterns. The heatmaps present scaled values of normalized counts from the RNA-seq data. (C) Expression patterns of 19 putative transcription factor genes picked and re-clustered from panel B.

We also determined that the expression patterns of some TF genes were consistent with the phenotypes. The expression levels of the orthologs of the bHLH TF genes *SPEECHLESS* (*SPCH*), *MUTE*, and *FAMA* were downregulated in developing submerged leaves, in which fewer stomata differentiated than in the aerial leaves (Figure 7A). In Arabidopsis, these genes are explicitly expressed in the guard cell lineage to promote stomatal differentiation (Ohashi-Ito and Bergmann, 2006; MacAlister et al., 2007; Pillitteri et al., 2008). Because these TFs are involved in stomatal development in many other land plants, including monocots and even mosses (Liu et al., 2009; Chater et al., 2016), it is likely they are also involved in stomatal development in *Callitriche* species. The stomatal lineage-associated expression of the genes encoding these TFs in several *Callitriche* species offers additional evidence of their functions in stomatal development (Doll et al., 2021). Among the other stomatal development-related genes, *EPIDERMAL PATTERNING FACTOR* (*EPF*) *1/2* orthologs and *TOO MANY MOUTHS* (*TMM*) genes were identified as DEGs in leaf primordia (Figure 7A). Additionally, the expression of the *EPFL9/STOMAGEN* ortholog was downregulated in submerged leaves. These genes were more highly expressed in the samples treated with AgNO_3_ and ABA than in the submerged control, although the differences were not always significant (Figure 7A).

A narrow, needle-like terete leaf form may result from the expansion of the abaxial region of a primordium (Waites and Hudson, 1995). This mechanism of narrowing was confirmed to be involved in the heterophylly of an aquatic plant species from the family Ranunculaceae (Kim et al., 2018). Thus, we also analyzed the expression of HD-ZIP III and KANADI TF genes, which determine the dorsiventral polarity (Figure 7A). Although some of the *KANADI* family genes were expressed at low levels in the submerged primordia, the overall expression profiles of the dorsiventral genes were not indicative of abaxialization.

### Candidate gene set for the regulation of heterophyll**y**

Finally, we compared transcriptome data to extract a set of genes that may be important for the differential development of *C*. *palustris* leaves. Of the 287 genes extracted from the comparisons of *C*. *palustris* experimental conditions, we excluded 22 genes that were orthologous to DEGs in *C*. *terrestris* because the consistent differential expression in *C*. *terrestris* and *C*. *palustris* implies that they may not be critical for differential leaf development (Figure 5L, M). Another 15 genes were also excluded because their expression levels were downregulated in the developing submerged leaves of *C*. *palustris*, but their orthologs were not expressed in *C*. *terrestris*, suggesting that the downregulated expression of these genes does not affect leaf formation. Thus, 250 genes, including 145 genes lacking an identified ortholog, were retained to form the candidate gene set for heterophylly regulation and differential development (Figure 7B; Supplemental Data Set 4). Of these genes, the coding regions were predicted for 208 genes.

The general importance of TFs in various developmental processes and environmental responses suggests that changes in TF gene expression levels are crucial for heterophylly. Therefore, we identified possible TF genes using PlantTFDB 4.0 (Jin et al., 2017) and via manual annotations. Among 80,437 predicted protein-coding genes, 2,489 encoded TFs. We identified 19 potential TF genes among the 208 candidate genes with predicted coding regions. The expression of five TF genes was upregulated in submerged leaves (Figure 7C). Of the 19 identified TF genes, 13 were developmental stage-specific DEGs. In addition to the TF genes, the candidate gene set included receptor-like protein (RLP) genes, such as a *CLAVATA1* (*CLV1*) ortholog and *BARELY ANY MERISTEM* (*BAM*) *1/2/3* homologs (Figure 7A, B), and 12 histone-like genes with expression levels that were upregulated in submerged primordia (Figure 7B; Supplemental Data Set 4). Moreover, a GO enrichment analysis revealed many of the candidate genes encoded transporters with increased abundance in submerged primordia as well as TFs (Figure 7B; Supplemental Data Set 5).

## Discussion

### Heterophylly in *Callitriche* and other aquatic plants

In this study, we focused on *Callitriche* species to analyze the plasticity of leaf development. Recent studies clarified the molecular basis of heterophylly in several distant lineages of aquatic plants. In *Rorippa aquatica* and *Hygrophila difformis*, the compound leaf forms of submerged plants are associated with the transcriptional activation of class I KNOX genes in leaves (Nakayama et al., 2014; Li et al., 2017). However, orthologs of these genes are not expressed in the simple leaves of *C*. *palustris* leaves under submerged conditions. Hence, these genes were not in our candidate gene set. An earlier study confirmed that abaxialization resulting from changes in the expression of genes related to dorsiventral polarity is important for heterophylly in *Ranunculus trichophyllus* (Kim et al., 2018). In contrast, our analysis of KANADI and HD-ZIP III TF gene expression in *C*. *palustris* leaves did not produce evidence of abaxialization (Figure 7A). These genes were also not in the candidate gene set. In *R. trichophyllus*, the activation of ethylene signaling and repression of ABA signaling contribute to the upregulated and downregulated expression of KANADI and HD-ZIP III genes, respectively. Thus, similar hormonal changes, but varying downstream gene expression, are presumed to occur in *C. palustris*. The differences in the mechanisms regulating heterophylly reflect the independent evolution of heterophylly in these plant species.

### Molecular background of dimorphic leaf formation in *C*. *palustris*

On the basis of transcriptome comparisons, we included 250 genes in the candidate gene set that regulates heterophylly in *C*. *palustris.* Although we currently do not know the development-related functions of these genes in *C*. *palustris*, some of the genes had expression patterns that are consistent with those in previous studies of genes in model organisms, including genes involved in stomatal development. We detected downregulated expression of *bHLH*, *EPF1/2*, and *TMM* genes in submerged leaves. In Arabidopsis, these genes are expressed almost exclusively in the stomatal lineage cells (Ohashi-Ito and Bergmann, 2006; MacAlister et al., 2007; Hara et al., 2007, 2009; Pillitteri et al., 2008), with only the *TMM* genes also expressed in the surrounding epidermal cells (Nadeau and Sack, 2002). Thus, the expression patterns for these genes in the current study may be due to a decrease in the number of stomatal lineage cells in submerged leaves. In addition to the genes expected to be expressed in the stomatal lineage cells, the expression levels of *EPFL9/STOMAGEN* orthologs were also downregulated in submerged primordia. In Arabidopsis, *EPFL9/STOMAGEN* is expressed in mesophyll cells and positively controls stomatal development in epidermal cells (Sugano et al., 2010). The external application of EPFL9*/*STOMAGEN leads to increased stomatal density, whereas knocking down *EPFL9/STOMAGEN* via RNA interference has the opposite effect (Sugano et al., 2010; Tanaka et al., 2013). Thus, the downregulated *EPFL9* expression potentially explains the decreased stomatal density in *C*. *palustris*. The suppression of *EPFL9/STOMAGEN* expression in submerged leaves was also observed in *R. trichophyllus*, likely because of activation of ethylene signaling and decreased ABA levels and the resulting inhibition of HD-ZIP III (Kim et al., 2018). In *C. palustris*, however, although the ABA content decreased in the submerged condition, HD-ZIP III gene expression was not repressed (Figure 7A). Thus, it is possible that stomatal control differs between *C. palustris* and *R. trichophyllus* leaves. Genetic analyses of stomatal regulation in *C. palustris*, including the involvement of ethylene signaling, are required to enable comparisons of the mechanisms.

Our candidate gene set included many genes encoding RLPs. These proteins are important for stress responses (e.g., defense against bacteria and viruses), but they also function as receptors for hormones and small peptide ligands, which influence various growth and developmental processes. An ortholog of *CLV1* and orphan *BAM*-like genes were also included in the candidate gene set (Figure 7A). The functions of the encoded proteins related to the regulation of the stem cell lineage in the shoot apical meristem are well known (DeYoung et al., 2006; Clark et al., 1997; DeYoung and Clark, 2008), but, in Arabidopsis, these genes are broadly expressed in the plant body, including leaves (Klepikova et al., 2016). For example, the *CLV1* promoter is active in leaf mesophyll cells (Kawade et al., 2013), although the effect of *CLV1* expression in leaves remains unreported. On the other hand, BAM genes are involved in xylem development in Arabidopsis (Qian et al., 2018). Therefore, the downregulated expression of the BAM orthologs in submerged *C. palustris* primordia may be associated with a decrease in the number of leaf veins (Koga et al., 2020). Although the changes in the expression of these receptor-like genes may be the result of changes in cellular state, rather than the drivers of cell differentiation, functionally characterizing these receptor-like genes may help elucidate the molecular basis of differential leaf development.

In the context of developmental biology, it is worthwhile to focus on TF genes in the candidate gene set. Because heterophylly of *C*. *palustris* is accompanied by significant changes in cell shape, it is likely that distinct differentiation processes mediate the formation of each leaf type. In Arabidopsis, some candidate TF gene homologs affect organ elongation. Both *OBP4L1* (Cluster-23900.0) and *OBP4L2* (Cluster-57510.0), which are Dof-type TF genes with downregulated expression in submerged primordia, are homologs of Arabidopsis *OBP4*, which is reportedly a negative regulator of cell growth (Xu et al., 2016; Rymen et al., 2017). Additionally, *PRE1A* (Cluster-74746.0) and *PRE1B* (Cluster-74746.1) expression levels were upregulated in submerged primordia. Their Arabidopsis homologs, *PACLOBUTRAZOL RESISTANCE* genes (*PRE1–6*), promote cell elongation in the hypocotyls, stems, and leaf petioles in a process controlled by GA signaling (Shin et al., 2019; Lee et al., 2006; Zhang et al., 2009). On the basis of the Arabidopsis phenotypes resulting from the disruption of these genes, it is unlikely that the expression-level changes in only one of these genes can substantially induce leaf blade elongations in *C. palustris*. We speculate that the coordinated effects of these genes and possibly other functionally unknown TF genes are responsible for the specific differentiation of elongated leaf cells in submerged leaf primordia. The molecular mechanism regulating heterophylly will be more precisely elucidated by studies focusing on these genes and how the hormones regulate their expression.

### Hormonal control of heterophylly

In a previous study on *C*. *heterophylla*, a GA application under aerial conditions induced the development of submerged-type leaves (Deschamp and Cooke, 1983). However, we were unable to replicate this result. Instead, in our scalable experimental system, the medium-cultured plants produced slightly elongated leaves in response to the GA_3_ application under aerial growth conditions. In these leaves, the cells were still highly lobed in the epidermis and had a relatively low aspect ratio in both layers (Supplemental Figures 5, 6), suggesting that GA signaling is insufficient to induce extensive cell elongation and the formation of submerged-type leaves. Thus, undesirable environmental factors may have affected the conditions that the soil-cultured plants were exposed to in the earlier study. Nevertheless, GA biosynthesis, as well as the subsequent activation of GA signaling, are likely required for the formation of submerged-type leaves. Notably, the reorientation of cMTs was associated with this phenomenon because GA signaling promotes cell elongation through the reorientation of cMTs and the cellulose microfibers (Mita and Shibaoka, 1984; Takeda and Shibaoka, 1981; Lloyd, 1991). Indeed, GA signaling was possibly more activated under submerged conditions than under aerial conditions, with upregulated *GA2OX* expression levels except in uniconazole P-treated primordia (Figure 6). Because our quantitative analysis of hormones failed to detect increases in GA contents, this activation may be caused by an undetectable increase in GA levels or increased sensitivity to GA (Figure 6). However, although GA signaling is required to promote cell elongation during the formation of submerged leaves in *C*. *palustris*, it alone seemed insufficient for modulating cMT orientations to the extent necessary to produce a submerged leaf. Considering uniconazole P was unable to completely block submerged-type cell formation, it is likely that GA signaling is not required for determining cellular fate, but is needed to promote general cell expansion regardless of the condition (Figure 8).

**Figure 8:**
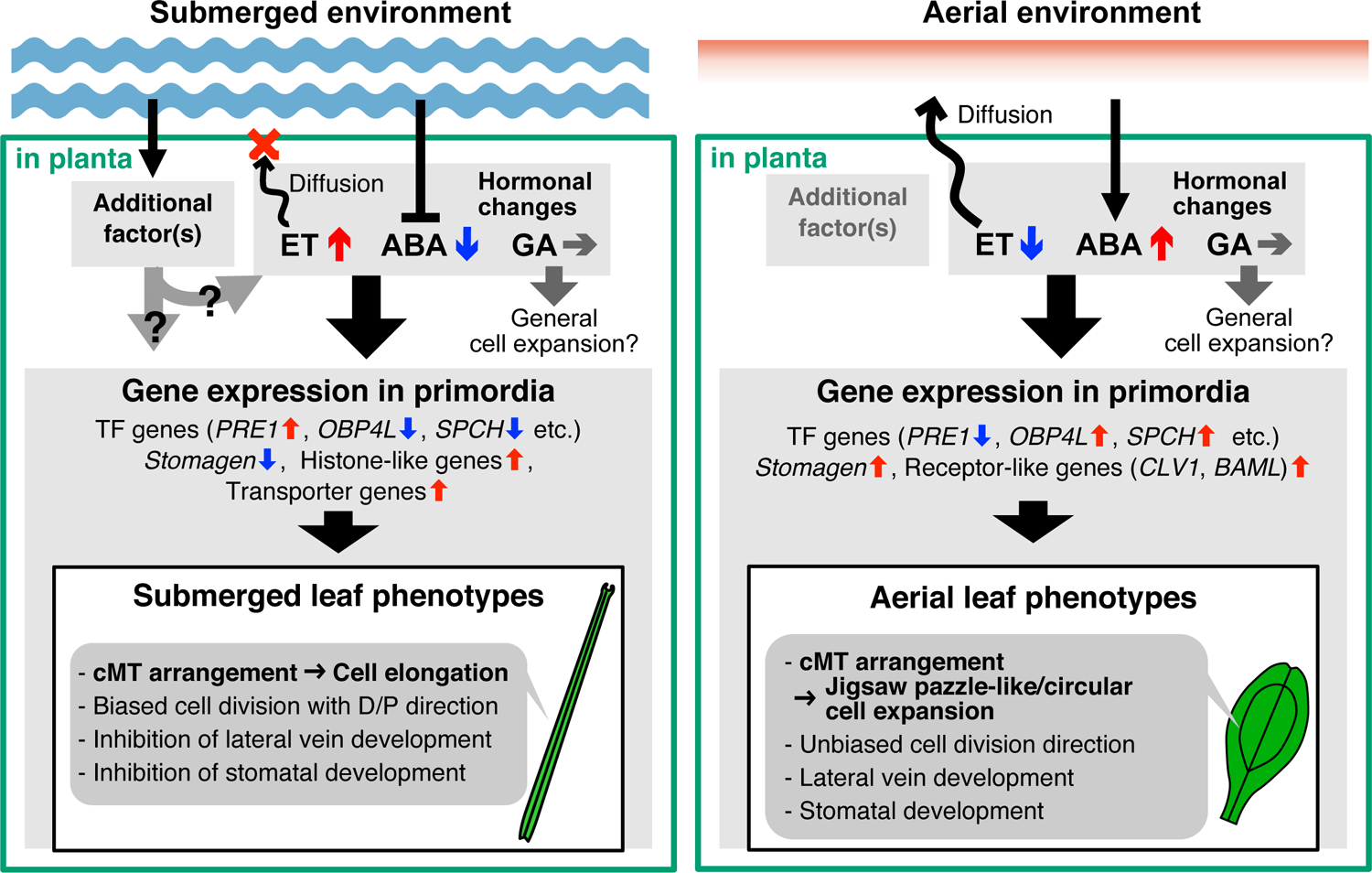
Schematic model of heterophylly in *C. palustris*. (A) In the submerged condition, ethylene signaling is activated, whereas ABA signaling is repressed. GA signaling may not necessarily be enhanced for submerged leaf formation. Either of the changes in hormone signaling alone is insufficient to induce the development of the submerged leaf phenotype. Additional factor(s), such as hypoxic stress, may be involved parallelly or additively with hormone signaling. The physiological changes induce gene expression changes in leaf primordia (< 500 µm long), after which leaf developmental processes associated with submerged-type cellular differentiation proceed. (B) In the aerial condition, while ethylene does not accumulate in the plant as it can diffuse outward, ABA production is promoted. Then, in the leaf primordia, many TF genes and some receptor-like genes, which are likely to be associated with the aerial leaf phenotype, are upregulated.

In some aquatic plants, ethylene positively regulates the formation of submerged leaves. For example, when *Ludwigia arcuate* (Onagraceae) is treated with ethylene gas or ACC under aerial growth conditions, the plants produce submerged-type leaves, indicating that ethylene signaling almost sufficiently induces submerged-type leaf development in this species (Kuwabara et al., 2003; Sato et al., 2008). Similarly, treating the aerial parts of *R. trichophyllus* and *Hygrophila defformis* plants with ethylene induces drastic changes in some leaf phenotypes, which are similar to those observed in submerged leaves to some extent (Kim et al., 2018; Li et al., 2017; Horiguchi et al., 2019). The involvement of ethylene is reasonable because, under submerged conditions, ethylene spontaneously accumulates in the plant body because of the limited gas exchange in water (Jackson, 1985). Indeed, substantial increases in ethylene levels have been reported for various plants growing under submerged conditions (Voesenek and Sasidharan, 2013) for a review). In *C*. *palustris*, an exposure to AgNO_3_, which inhibits the perception of ethylene, induced the production of aerial-type leaves under submerged conditions, which suggests that ethylene signaling must be activated for the formation of submerged leaves. However, this mechanism seems to be insufficient for the formation of submerged leaves in *C*. *palustris* because the application of ethylene or ACC did not induce the formation of submerged leaves. Regarding cellular morphology, the activation of ethylene had minimal effects and did not lead to submerged leaf phenotypes (Figure 3; Supplemental Figure 5). This is consistent with the inhibition of submerged leaf formation by uniconazole P under submerged conditions, in which ethylene should spontaneously accumulate. This implies that the formation of submerged leaves is a GA-dependent process. Nevertheless, we were unable to induce the formation of submerged leaves through the combined application of GA_3_ and ACC. The inhibition of ethylene signaling resulted in aerial-like cellular morphologies even with GA_3_ (Figure 2), implying that ethylene signaling has a greater role than GA signaling in submerged-type cell differentiation. Collectively, these results indicate that *C. palustris* requires ethylene signaling for submerged-type cell differentiation, but it is nearly insensitive to ethylene under aerial conditions. Additional factors associated with submergence are likely needed for ethylene signaling to trigger the formation of submerged-type leaves.

In the current study, the application of ACC or ethylene, even with GA_3_, induced seemingly normal stress responses commonly observed in terrestrial plants (e.g., strong growth inhibition and chlorosis) (Supplemental Figure 3). This ethylene response was inconsistent with the healthy growth of the submerged plants. Interestingly, in experiments conducted by Musgrave *et al*. (1972), applying ethylene gas to the floating rosettes of *Callitriche platycarpa* plants enhanced stem elongation, but the leaf forms were not thoroughly examined. Additionally, an effective amount of ethylene for promoting stem elongation accumulated in the submerged plants. These results suggest that at least in the floating shoots, where the basal part of the plant is submerged, the plant reacts to ethylene with an elongation response, unlike in the fully aerial condition (Musgrave et al., 1972). The mechanism mediating the contradictory effects of ethylene signaling under submerged and aerial conditions remains unclear, because the genetic basis of ethylene signaling in *C*. *palustris* has not been characterized. Thus, molecular investigations of the ethylene responses are required to fully elucidate the heterophylly of *C*. *palustris*.

Abscisic acid is also known as an inhibitory hormone of submerged leaf formation in broad aquatic plants, including ferns (Wanke 2011). Regarding *Callitriche* species, an ABA treatment under submerged conditions reportedly leads to the formation of aerial-like leaves (Deschamp and Cooke 1983). In the present study, we proved that the ABA content decreases substantially in the submerged shoot, possibly because of the downregulated expression of NCED-encoding genes in mature leaves. Our leaf and cellular measurements confirmed that the application of exogenous ABA under submerged conditions resulted in leaves that were morphologically identical to aerial leaves, indicating that ABA levels must decrease before submerged leaves can form. However, because the ABA content also decreased in the AgNO_3_- and uniconazole P-treated *C palustris* shoots, a decrease in ABA levels alone is insufficient for inducing submerged leaf formation. Notably, the ABA receptor orthologs *PYR1* and *PYL4/5/6* were differentially expressed between the heterophyllous leaf primordia (Figure 6B). Moreover, three *C. palustris*-specific *PYL4/5/6*-like genes were included in the candidate gene set (Figure 7B, Supplemental Data Set 4). The downregulated expression of *CpPYR1* in submerged leaves can be explained by reductions in the number of guard cells and the amount of vascular tissue because *PYR1* gene is specifically expressed in these tissues in Arabidopsis (Gonzalez-Guzman et al., 2012). The *PYL4* expression levels were markedly upregulated in the submerged primordia of both *C. palustris* and *C. terrestris*, implying changes in the expression of these genes are not associated with the morphological changes. However, CpPYL4 may contribute to physiological changes in the submerged form by modulating the sensitivity to ABA or inducing differential responses downstream of the ABA signaling pathway.

### Requirement of additional factors for the formation of submerged leaves

The results of our hormone perturbation experiments suggest that factor(s) other than ethylene and GA are also required to induce the development of submerged leaves under aerial conditions. Our results of hormonal measurements suggested a reduction of ABA amount is one of the requirements. Because the submerged condition is a drastically different environment from the aerial condition, various factors can be assumed, such as changes in light quality, photoperiod, and temperature. These factors have previously been demonstrated to induce differential leaf formation in other aquatic plants (Wells and Pigliucci, 2000). One such factor to be considered may be high turgor pressure under submerged conditions (Deschamp and Cooke, 1983). Low-oxygen stress is another potential factor. Indeed, in *R. trichophyllus*, hypoxia can partly induce the development of submerged leaf phenotypes under aerial conditions (Kim et al., 2018).

Group VII ERFs, which are activated in Arabidopsis by hypoxic conditions, influence the morphological responses of submerged rice plants (Hattori et al., 2009; Fukao et al., 2006; Xu et al., 2006), and may affect the viability of *Rumex palustris* and *Rorippa* species under submerged conditions (Licausi et al., 2011; VEEN et al., 2014). The expression patterns of the group VII ERFs imply the hypoxic response is probably activated in both species under submerged conditions, with the underlying mechanism controlled by ethylene signaling in *C. palustris* (Figure 7A). However, the upregulated expression of the genes encoding the group VII ERFs was not correlated with leaf shapes. Thus, this pathway is insufficient for the formation of submerged leaves in *C*. *palustris*, or it may not be involved at all.

Considered together, our findings confirm that the heterophylly of *C. palustris* is controlled by a complex mechanism involving phytohormones and possibly additional factors (Figure 8A). Expression-level changes in relatively few genes, including diverse TF genes, might be primarily involved in modulating cellular states for submerged-type leaf phenotypes (e.g., rearrangement of cMT orientation and subsequent cell elongation and inhibition of stomatal development). The considerable cell elongation in submerged leaves requires GA signaling, but GA contents do not increase following submergence. Ethylene signaling is also required for the cell elongation in submerged leaves. In contrast, the abundance of ABA, which promotes aerial leaf formation, must decrease prior to submerged leaf formation (Figure 8A). In the aerial condition, the ABA signaling is activated, but the ethylene signaling is not as it can volatilize to the atmosphere immediately, leading to aerial-type leaf phenotypes (e.g., jigsaw puzzle-like or circular cell expansion, and stomatal and vein development; Figure 8B). These hormones are similarly involved in the heterophylly of amphibious plants from different lineages, but the changes to genetic architectures downstream of these hormone signals likely differ among the lineages. In *C. palustris*, the KNOX genes (Nakayama et al 2014, Li et al, 2017) as well as the ethylene–KANADI and ABA–HD-ZIP III modules (Kim et al., 2018) may not be involved in heterophylly. It is also notable that *C. palustris* is only slightly responsive to hormone applications under aerial conditions. In many amphibious plants, hormone applications drastically alter leaf phenotypes, although not always resulting in a completely submerged leaf form, suggesting that the extent of the contributions of the same hormones to heterophylly varies among lineages. These observations shed light on the variety of evolutionary trajectories of plant adaptations to aquatic environments.

## Materials and Methods

### Plant culture

*Callitriche palustris* L. and *Callitriche terrestris* Raf. plants were originally collected in Hakuba, Nagano, Japan (NH1 strain), and Nishinomiya, Hyogo, Japan, respectively. The plant culture system used in this study was recently described by Koga *et al*. (2020). Briefly, the plant material from a sterile culture was transplanted onto solid half-strength MS medium after the visible shoot tips were removed. A culture jar was filled with 200 mL sterile distilled water to produce submerged growth conditions. For the chemical treatments under submerged conditions, the chemical to be tested was diluted in 200 mL sterile distilled water. The specific concentrations for each experiment are indicated in the corresponding figures and/or figure legends. For chemical treatments under aerial conditions, the chemical was added to the medium in advance or sprayed to the shoots every 2 or 3 days. For the chemical treatments under aerial conditions, the chemical was added to the medium in advance or was applied by spraying the shoots every 2 or 3 days. Regarding the ethylene gas treatment, the plants were transplanted into airtight flasks containing the growth medium, after which they were exposed to ethylene gas every two days.

### Measurement of leaves and cells

Before measuring the leaves and cells, the plants were treated with a formalin–acetic acid–alcohol fixative. The leaves from the fixed shoots were dissected and then examined. Images of the leaves were captured with a scanner or a camera attached to a microscope. The length of the main vein was recorded as the leaf length, whereas the length of the widest line that vertically crossed the main vein was recorded as the leaf width. ImageJ/Fiji software was used to analyze the scanned images to measure each index. The examined leaves were collected from two or three sibling plants as biological replicates for each experiment. For cellular observations, the fixed leaves were cleared in a chloral hydrate solution (Tsuge et al., 1996). The leaf cells were observed using a DM4500 differential interference contrast microscope (Leica Microsystems, Wetzlar, Germany). We obtained cellular images from 4–8 leaves collected from two or three sibling plants as biological replicates. Using ImageJ/Fiji, we manually traced the contours of 30 pavement cells and 30 subepidermal palisade cells per leaf. The analyzed cells were randomly selected, but we avoided choosing pavement cells adjacent to guard cells and hair cells because their shapes were distorted. We then examined the cell size and other shape-related indices using the “Area,” “Shape descriptors,” and “Perimeter” measurement functions of ImageJ/Fiji (Supplemental Figure 1; see ImageJ documents for more details). The cellular length and width were measured as the long and short side lengths of the bounding smallest rectangle, respectively (Supplemental Figure 1). Statistical analyses and visualizations were performed using the following packages of the R software (version 4.0.3): multcomp (version 1.4-16) (Hothorn et al., 2008), tidyverse (version 1.3.0) (Wickham et al., 2019), and ggforce (version 0.3.3; https://CRAN.R-project.org/package=ggforce), and ggfortify (version 0.4.11) (Tang et al., 2016).

### Immunofluorescence

The immunofluorescence of microtubules was analyzed as previously described (Pasternak et al., 2015), with slight modifications. Briefly, the leaves were fixed in 2% PFA/0.5% glutaraldehyde prepared in a microtubule-stabilizing buffer, after which the central region was dissected. After permeabilizing with methanol, cell wall enzymes, and a 3% (v/v) IGEPAL/10% (v/v) DMSO solution, the samples were incubated with a 1:1,000 primary antibody solution (T5168; Sigma-Aldrich, St. Louis, MO, USA) and then with a 1:500 Alexa Fluor 488-conjugated secondary antibody solution (A-11017; Thermo Fisher Scientific, Waltham, MA, USA). The stained samples were observed using an FV10 confocal microscope (Olympus, Tokyo, Japan).

### RNA-seq

For the transcriptome analyses, we cultured shoots from individual *C*. *palustris* and *C*. *terrestris* plants grown under aerial or submerged conditions. After culturing for three weeks, we extracted RNA from whole plants using the RNeasy Plant Mini Kit (Qiagen, Hilden, Germany) and the RNase-free DNase Set (Qiagen). Sequencing libraries were prepared using the TruSeq Stranded mRNA Library Prep Kit (Illumina, Inc., San Diego, CA, USA), with the protocol optimized for a 300–400 bp insert size. We sequenced the libraries on the HiSeq 1500 platform (Illumina) using the rapid-run mode to generate 150-bp paired-end reads. Additionally, for the quantitative analysis of RNA, we cultured shoots from three siblings grown under aerial conditions, submerged conditions, and submerged conditions with 10^−6^ M AgNO_3,_ 10^−7^ M uniconazole P, or 10^−7^ M ABA. We extracted total RNA from 10 leaves that were shorter than 500 µm and from 4–6 mature leaves using the RNeasy Micro Kit (Qiagen) and the RNeasy Plant Mini Kit (Qiagen), respectively, along with the RNase-free DNase Set (Qiagen). We constructed sequencing libraries with the KAPA Stranded mRNA-seq Kit (KAPA Biosystems, Inc., Wilmington, MA, USA), after which they were sequenced on the HiSeq 1500 platform (Illumina) using the rapid-run mode to obtain 75-bp single-read sequences. The raw reads were deposited in the DDBJ Sequence Read Archive (DRA) (Supplemental Data Set 2).

### Transcriptome assembly and annotation

The paired-end reads for *C*. *palustris* or *C*. *terrestris* whole plants were trimmed based on quality using Trimmomatic (version 0.36) (minimum length: 32) (Bolger et al., 2014) and assembled with Trinity (version 2.2.0) with an *in silico* normalization step (Grabherr et al., 2011). The contigs derived from rRNA were identified using RNAmmer (version 1.2.1) (Lagesen et al., 2007) and removed from the assembly. Additionally, contigs with the best matches (e-value < 1E-50) to sequences from non-Viridiplantae species (mostly bacteria, fungi, and human) following a BLASTn search of the nt database were considered as possible contaminants and eliminated. The single open reading frame (ORF) of each transcript was predicted using TransDecoder (version 3.0.0) (https://github.com/TransDecoder/TransDecoder). The assemblies included many contigs putatively transcribed from a single gene. Because the clustering intrinsically executed by Trinity did not work properly for the tetraploid *C*. *palustris* transcriptome, we clustered contigs into putative genes using Corset (version 1.07) (Davidson and Oshlack, 2014). To select a representative isoform of each putative gene, the expression levels of isoforms were quantified by mapping the reads onto the assembly using Bowtie (version 1.2) and RSEM (version 2.3.0) (Li and Dewey, 2011; Langmead et al., 2009). Representative isoforms were selected based on the following criteria: having an ORF, BLASTx match to a sequence in the UniProt database, high expression level, and a relatively long transcript among the isoforms for the gene. Next, we qualitatively analyzed the assembly using BUSCO (version 4.0.1) with the Eudicot data set (Simão et al., 2015). Briefly, genes were annotated and GO terms were assigned based on the BLASTx best match between representative genes and Arabidopsis protein sequences (TAIR10). Orthologs were detected with OrthoFinder (version 2.1.2) (Emms and Kelly, 2015), using the predicted protein sequences of the species, and the protein sequences from Arabidopsis, tomato (ITAG2.4), and *Mimulus guttatus* (version 2.0 in Phytozome 11.0) as outgroups. We subsequently identified genes with one-to-one or multi-to-one relationships between tetraploid *C*. *palustris* and diploid *C*. *terrestris* as comparable orthologs.

### Comparative expression analyses

The reads obtained for the leaves were trimmed based on quality using Trimmomatic (version 0.36), after which they were counted by RSEM by mapping them onto the transcriptome assembly with Bowtie. Genes expressed at low levels (TPM < 1 in all samples) were omitted from further analyses. After a normalization using the TCC package (version 1.20.0) (Sun et al., 2013), the DEGs between certain sample pairs were detected via multiple comparisons using edgeR (version 3.22.1) (Robinson and Oshlack, 2010), with the following criteria: FDR < 0.05 and expression-level log_2_(fold-change) > 1. The GO term enrichment analyses were performed using R software (version 4.0.3). Specifically, we applied a hypergeometric distribution and an FDR < 0.05 calculated by a BH correction. Rare ontologies (< 5 in the transcriptome) were omitted from the analyses. The data were visualized using the following packages of the R software: tidyverse (version 1.3.0) (Wickham et al., 2019), ggfortify (version 0.4.11) (Tang et al., 2016), factoextra (version 1.0.7; https://CRAN.R-project.org/package=factoextra), UpSetR (version 1.4.0) (Conway et al., 2017), Vennerable (version 3.1.0.9000; https://github.com/js229/Vennerable), and ComplexHeatmap (version 2.5.3) (Gu et al., 2016).

### Hormonal amount measurement

The shoot tips with developing and almost mature leaves were collected from plants cultured under the same conditions as the plants used for the RNA-seq experiments. Hormones were extracted and semi-purified from the frozen samples as previously described (Kojima and Sakakibara, 2012; Kojima et al., 2009).

The GA, ABA, salicylic acid, jasmonic acid, and auxin contents were quantified using an ultra-high performance liquid chromatography (UHPLC)-ESI quadrupole-orbitrap mass spectrometer (UHPLC/Q-Exactive; Thermo Fisher Scientific, USA) with an ODS column (AQUITY UPLC HSS T3, 1.8 µm, 2.1 × 100 mm; Waters, Milford, MA, USA) as previously described (Kojima and Sakakibara, 2012; Shinozaki et al., 2015). Cytokinins were quantified using an ultra-performance liquid chromatography (UPLC)-electrospray interface (ESI) tandem quadrupole mass spectrometer (qMS/MS) (AQUITY UPLC System/Xevo-TQS; Waters) with an ODS column (AQUITY UPLC HSS T3, 1.8 µm, 2.1 × 100 mm; Waters) as previously described (Kojima et al., 2009).

### Accession numbers

Sequence data used in this article can be found in the DDBJ DRA with the accession numbers listed in Supplemental table Data Set 2.

## Acknowledgments

We are grateful to H. Fukuda and Y. Kondo at the University of Tokyo for kindly allowing us to use their confocal microscope. Computational resources were provided by the Data Integration and Analysis Facility, National Institute for Basic Biology. We thank Edanz Group (https://en-author-services.edanz.com/ac) for editing a draft of this manuscript. This work was supported by a Grant-in-Aid for Research Activity start-up, a Grant-in-Aid for Early-Career Scientists, and a Grant-in-Aid for Scientific Research on Innovative Areas (Grant Nos. JP16H06733, 20K15816, JP25113002, and JP19H05672).

## Author contributions

H.K. and H.T. designed the research; H.K., M.K., Y.T, and H.S. conducted the experiments; H.K. analyzed the data; and all authors contributed to writing the manuscript.

## References

1. Allsopp, A. (1967). Heteroblastic Development in Vascular Plants. Advances in Morphogenesis 6: 127–171.

2. Arber, A. (1920). Water plants: a study of aquatic angiosperms (Cambridge University Press: Cambridge).

3. Bolger, A.M., Lohse, M., and Usadel, B. (2014). Trimmomatic: a flexible trimmer for Illumina sequence data. Bioinformatics 30: 2114–2120.

4. Bradshaw, A.D. (1965). Evolutionary significance of phenotypic plasticity in plants. Adv Genet 13: 115–155.

5. Chater, C.C. et al. (2016). Origin and function of stomata in the moss *Physcomitrella patens*. Nat Plants 2: 16179.

6. Clark, S.E., Williams, R.W., and Meyerowitz, E.M. (1997). The CLAVATA1Gene Encodes a Putative Receptor Kinase That Controls Shoot and Floral Meristem Size in Arabidopsis. Cell 89: 575–585.

7. Conway, J.R., Lex, A., and Gehlenborg, N. (2017). UpSetR: an R package for the visualization of intersecting sets and their properties. Bioinformatics 33: 2938–2940.

8. Cook, C.D.K. (1999). The number and kinds of embryo-bearing plants which have become aquatic: a survey. Perspect Plant Ecol Syst 2: 79–102.

9. Cox, M.C.H., Benschop, J.J., Vreeburg, R.A.M., Wagemaker, C.A.M., Moritz, T., Peeters, A.J.M., and Voesenek, L.A.C.J. (2004). The Roles of Ethylene, Auxin, Abscisic Acid, and Gibberellin in the Hyponastic Growth of Submerged *Rumex palustris* Petioles. Plant Physiol 136: 2948–2960.

10. Davidson, N.M. and Oshlack, A. (2014). Corset: enabling differential gene expression analysis for de novo assembled transcriptomes. Genome Biol 15: 410.

11. Deschamp, P.A. and Cooke, T.J. (1984). Causal mechanisms of leaf dimorphism in the aquatic angiosperm *Callitriche heterophylla*. Am J Bot 71: 319–329.

12. Deschamp, P.A. and Cooke, T.J. (1983). Leaf Dimorphism in Aquatic Angiosperms: Significance of Turgor Pressure and Cell Expansion. Science 219: 505–507.

13. Deschamp, P.A. and Cooke, T.J. (1985). Leaf Dimorphism in the Aquatic Angiosperm *Callitriche heterophylla*. Am J Bot 72: 1377.

14. DeYoung, B.J., Bickle, K.L., Schrage, K.J., Muskett, P., Patel, K., and Clark, S.E. (2006). The CLAVATA1 related BAM1, BAM2 and BAM3 receptor kinase like proteins are required for meristem function in Arabidopsis. Plant J 45: 1–16.

15. DeYoung, B.J. and Clark, S.E. (2008). BAM Receptors Regulate Stem Cell Specification and Organ Development Through Complex Interactions With CLAVATA Signaling. Genetics 180: 895–904.

16. Doll, Y., Koga, H., and Tsukaya, H. (2021). The diversity of stomatal development regulation in *Callitriche* is related to the intrageneric diversity in lifestyles. Proc Natl Acad Sci USA 118: e2026351118.

17. Du, Z.-Y. and Wang, Q.-F. (2014). Correlations of Life Form, Pollination Mode and Sexual System in Aquatic Angiosperms. Plos One 9: e115653.

18. Emms, D.M. and Kelly, S. (2015). OrthoFinder: solving fundamental biases in whole genome comparisons dramatically improves orthogroup inference accuracy. Genome Biol 16: 157.

19. Erbar, C. and Leins, P. (2004). Callitrichaceae. In, pp. 50–56.

20. Fassett, N.C. (1951). *Callitriche* in the new world. Rhodora 53: 161.

21. Fukao, T., Xu, K., Ronald, P.C., and Bailey-Serres, J. (2006). A Variable Cluster of Ethylene Response Factor–Like Genes Regulates Metabolic and Developmental Acclimation Responses to Submergence in Rice. Plant Cell 18: 2021–2034.

22. Gonzalez-Guzman, M., Pizzio, G.A., Antoni, R., Vera-Sirera, F., Merilo, E., Bassel, G.W., Fernández, M.A., Holdsworth, M.J., Perez-Amador, M.A., Kollist, H., and Rodriguez, P.L. (2012). Arabidopsis PYR/PYL/RCAR Receptors Play a Major Role in Quantitative Regulation of Stomatal Aperture and Transcriptional Response to Abscisic Acid. Plant Cell 24: 2483–2496.

23. Grabherr, M.G. et al. (2011). Full-length transcriptome assembly from RNA-Seq data without a reference genome. Nat Biotechnol 29: 644–652.

24. Green, P.B. (1980). Organogenesis-A Biophysical View. Ann Rev Plant Physio 31: 51–82.

25. Gu, Z., Eils, R., and Schlesner, M. (2016). Complex heatmaps reveal patterns and correlations in multidimensional genomic data. Bioinformatics 32: 2847–2849.

26. Gunning, B.E. and Hardham, A.R. (1982). Microtubules. Annu Rev Plant Physiol 33: 651– 698.

27. Hara, K., Kajita, R., Torii, K.U., Bergmann, D.C., and Kakimoto, T. (2007). The secretory peptide gene EPF1 enforces the stomatal one-cell-spacing rule. Gene Dev 21: 1720–1725.

28. Hara, K., Yokoo, T., Kajita, R., Onishi, T., Yahata, S., Peterson, K.M., Torii, K.U., and Kakimoto, T. (2009). Epidermal Cell Density is Autoregulated via a Secretory Peptide, EPIDERMAL PATTERNING FACTOR 2 in Arabidopsis Leaves. Plant Cell Physiol 50: 1019–1031.

29. Hattori, Y. et al. (2009). The ethylene response factors SNORKEL1 and SNORKEL2 allow rice to adapt to deep water. Nature 460: 1026–1030.

30. Horiguchi, G., Nemoto, K., Yokoyama, T., and Hirotsu, N. (2019). Photosynthetic acclimation of terrestrial and submerged leaves in the amphibious plant *Hygrophila difformis*. Aob Plants 11: plz009.

31. Hothorn, T., Bretz, F., and Westfall, P. (2008). Simultaneous Inference in General Parametric Models. Biometrical J 50: 346–363.

32. Ito, Y., Tanaka, N., Barfod, A.S., Kaul, R.B., Muasya, A.M., Garcia-Murillo, P., Vere, N.D., Duyfjes, B.E.E., and Albach, D.C. (2017). From terrestrial to aquatic habitats and back again: molecular insights into the evolution and phylogeny of *Callitriche* (Plantaginaceae). Bot J Linn Soc 184: 46–58.

33. Jackson, M.B. (1985). Ethylene and Responses of Plants to Soil Waterlogging and Submergence. Annu Rev Plant Phys 36: 145–174.

34. Jackson, M.B. (2008). Ethylene-promoted Elongation: an Adaptation to Submergence Stress. Ann Bot 101: 229–248.

35. Jin, J., Tian, F., Yang, D.-C., Meng, Y.-Q., Kong, L., Luo, J., and Gao, G. (2017). PlantTFDB 4.0: toward a central hub for transcription factors and regulatory interactions in plants. Nucleic Acids Res 45: D1040–D1045.

36. Jones, H. (1955). Further studies on heterophylly in *Callitriche intermedia*: leaf development and experimental induction of ovate leaves. Ann Bot 19: 370–388.

37. Jones, H. (1952). Variation in Leaf Form in *Callitriche intermedia*. Nature 170: 848–849.

38. Kawade, K., Horiguchi, G., Usami, T., Hirai, M.Y., and Tsukaya, H. (2013). ANGUSTIFOLIA3 Signaling Coordinates Proliferation between Clonally Distinct Cells in Leaves. Curr Biol 23: 788–792.

39. Kim, J., Joo, Y., Kyung, J., Jeon, M., Park, J.Y., Lee, H.G., Chung, D.S., Lee, E., and Lee, I. (2018). A molecular basis behind heterophylly in an amphibious plant, *Ranunculus trichophyllus*. Plos Genet 14: e1007208.

40. Klepikova, A.V., Kasianov, A.S., Gerasimov, E.S., Logacheva, M.D., and Penin, A.A. (2016). A high resolution map of the *Arabidopsis thaliana* developmental transcriptome based on RNA seq profiling. Plant J 88: 1058–1070.

41. Koga, H., Doll, Y., Hashimoto, K., Toyooka, K., and Tsukaya, H. (2020). Dimorphic Leaf Development of the Aquatic Plant *Callitriche palustris* L. Through Differential Cell Division and Expansion. Front Plant Sci 11: 269.

42. Kojima, M., Kamada-Nobusada, T., Komatsu, H., Takei, K., Kuroha, T., Mizutani, M., Ashikari, M., Ueguchi-Tanaka, M., Matsuoka, M., Suzuki, K., and Sakakibara, H. (2009). Highly Sensitive and High-Throughput Analysis of Plant Hormones Using MS-Probe Modification and Liquid Chromatography–Tandem Mass Spectrometry: An Application for Hormone Profiling in *Oryza sativa*. Plant Cell Physiol 50: 1201–1214.

43. Kojima, M. and Sakakibara, H. (2012). High-Throughput Phenotyping in Plants, Methods and Protocols. Methods Mol Biology 918: 151–164.

44. Kuwabara, A., Ikegami, K., Koshiba, T., and Nagata, T. (2003). Effects of ethylene and abscisic acid upon heterophylly in *Ludwigia arcuata* (Onagraceae). Planta 217: 880–887.

45. Kuwabara, A., Tsukaya, H., and Nagata, T. (2001). Identification of Factors that Cause Heterophylly in *Ludwigia arcuata* Walt. (Onagraceae). Plant Biol 3: 98–105.

46. Lagesen, K., Hallin, P., Rødland, E.A., Stærfeldt, H.-H., Rognes, T., and Ussery, D.W. (2007). RNAmmer: consistent and rapid annotation of ribosomal RNA genes. Nucleic Acids Res 35: 3100–3108.

47. Langmead, B., Trapnell, C., Pop, M., and Salzberg, S.L. (2009). Ultrafast and memory-efficient alignment of short DNA sequences to the human genome. Genome Biol 10: R25.

48. Lee, S., Lee, S., Yang, K.-Y., Kim, Y.-M., Park, S.-Y., Kim, S.Y., and Soh, M.-S. (2006). Overexpression of PRE1 and its Homologous Genes Activates Gibberellin-dependent Responses in *Arabidopsis thaliana*. Plant Cell Physiol 47: 591–600.

49. Li, B. and Dewey, C.N. (2011). RSEM: accurate transcript quantification from RNA-Seq data with or without a reference genome. Bmc Bioinformatics 12: 323.

50. Li, G., Hu, S., Hou, H., and Kimura, S. (2019). Heterophylly: Phenotypic Plasticity of Leaf Shape in Aquatic and Amphibious Plants. Plants 8: 420.

51. Li, G., Hu, S., Yang, J., Schultz, E.A., Clarke, K., and Hou, H. (2017). Water-Wisteria as an ideal plant to study heterophylly in higher aquatic plants. Plant Cell Rep 36: 1225–1236.

52. Licausi, F., Dongen, J.T.V., Giuntoli, B., Novi, G., Santaniello, A., Geigenberger, P., and Perata, P. (2010). HRE1 and HRE2, two hypoxia inducible ethylene response factors, affect anaerobic responses in *Arabidopsis thaliana*. Plant J 62: 302–315.

53. Licausi, F., Kosmacz, M., Weits, D.A., Giuntoli, B., Giorgi, F.M., Voesenek, L.A.C.J., Perata, P., and van Dongen, J.T. (2011). Oxygen sensing in plants is mediated by an N-end rule pathway for protein destabilization. Nature 479: 419–422.

54. Liu, T., Ohashi-Ito, K., and Bergmann, D.C. (2009). Orthologs of *Arabidopsis thaliana* stomatal bHLH genes and regulation of stomatal development in grasses. Development 136: 2265–2276.

55. Lloyd, C. (1991). The cytoskeletal basis of plant growth and form.

56. MacAlister, C.A., Ohashi-Ito, K., and Bergmann, D.C. (2007). Transcription factor control of asymmetric cell divisions that establish the stomatal lineage. Nature 445: 537–540.

57. Mita, T. and Shibaoka, H. (1984). Gibberellin stabilizes microtubules in onion leaf sheath cells. Protoplasma 119: 100–109.

58. Musgrave, A., Jackson, M.B., and Ling, E. (1972). *Callitriche* stem elongation is controlled by ethylene and gibberellin. Nat New Biol 238: 93.

59. Nadeau, J.A. and Sack, F.D. (2002). Control of Stomatal Distribution on the *Arabidopsis* Leaf Surface. Science 296: 1697–1700.

60. Nakayama, H., Nakayama, N., Seiki, S., Kojima, M., Sakakibara, H., Sinha, N., and Kimura, S. (2014). Regulation of the KNOX-GA Gene Module Induces Heterophyllic Alteration in North American Lake Cress. Plant Cell 26: 4733–4748.

61. Nakayama, H., Sinha, N.R., and Kimura, S. (2017). How Do Plants and Phytohormones Accomplish Heterophylly, Leaf Phenotypic Plasticity, in Response to Environmental Cues. Front Plant Sci 8: 1717.

62. Ohashi-Ito, K. and Bergmann, D.C. (2006). Arabidopsis FAMA Controls the Final Proliferation/Differentiation Switch during Stomatal Development. Plant Cell 18: 2493–2505.

63. Pasternak, T., Tietz, O., Rapp, K., Begheldo, M., Nitschke, R., Ruperti, B., and Palme, K. (2015). Protocol: an improved and universal procedure for whole-mount immunolocalization in plants. Plant Methods 11: 50.

64. Philbrick, C. (1994). Chromosome counts for *Callitriche* (Callitrichaceae) in North America. Rhodora 96: 383–386.

65. Philbrick, C.T. and Jansen, R.K. (1991). Phylogenetic Studies of North American *Callitriche* (Callitrichaceae) Using Chloroplast DNA Restriction Fragment Analysis. Syst Bot 16: 478.

66. Philbrick, C.T. and Les, D.H. (2000). Phylogenetic studies in *Callitriche*: implications for interpretation of ecological, karyological and pollination system evolution. Aquat Bot 68: 123– 141.

67. Pillitteri, L.J., Bogenschutz, N.L., and Torii, K.U. (2008). The bHLH Protein, MUTE, Controls Differentiation of Stomata and the Hydathode Pore in Arabidopsis. Plant Cell Physiol 49: 934–943.

68. Prančl, J., Kaplan, Z., Trávníček, P., and Jarolímová, V. (2014). Genome Size as a Key to Evolutionary Complex Aquatic Plants: Polyploidy and Hybridization in *Callitriche* (Plantaginaceae). Plos One 9: e105997.

69. Preston, R.D. (1974). The physical biology of plant cell walls (Chapman & Hall).

70. Qian, P., Song, W., Yokoo, T., Minobe, A., Wang, G., Ishida, T., Sawa, S., Chai, J., and Kakimoto, T. (2018). The CLE9/10 secretory peptide regulates stomatal and vascular development through distinct receptors. Nat Plants 4: 1071–1081.

71. Robinson, M.D. and Oshlack, A. (2010). A scaling normalization method for differential expression analysis of RNA-seq data. Genome Biol 11: R25.

72. Rymen, B., Kawamura, A., Schäfer, S., Breuer, C., Iwase, A., Shibata, M., Ikeda, M., Mitsuda, N., Koncz, C., Ohme-Takagi, M., Matsui, M., and Sugimoto, K. (2017). ABA Suppresses Root Hair Growth via the OBP4 Transcriptional Regulator. Plant Physiol 173: 1750– 1762.

73. Sato, M., Tsutsumi, M., Ohtsubo, A., Nishii, K., Kuwabara, A., and Nagata, T. (2008). Temperature-dependent changes of cell shape during heterophyllous leaf formation in *Ludwigia arcuata* (Onagraceae). Planta 228: 27–36.

74. Schenck, H. (1887). Vergleichende anatomie der submersen gewächse. Bibl bot 1: 1–67.

75. Sculthorpe, C.D. (1967). The biology of aquatic vascular plants (Edward Arnold: London).

76. Shin, K., Lee, I., Kim, E., Park, S., Soh, M.-S., and Lee, S. (2019). PACLOBUTRAZOL-RESISTANCE Gene Family Regulates Floral Organ Growth with Unequal Genetic Redundancy in *Arabidopsis thaliana*. Int J Mol Sci 20: 869.

77. Simão, F.A., Waterhouse, R.M., Ioannidis, P., Kriventseva, E.V., and Zdobnov, E.M. (2015). BUSCO: assessing genome assembly and annotation completeness with single-copy orthologs. Bioinformatics 31: 3210–3212.

78. Solano, R., Stepanova, A., Chao, Q., and Ecker, J.R. (1998). Nuclear events in ethylene signaling: a transcriptional cascade mediated by ETHYLENE-INSENSITIVE3 and ETHYLENE-RESPONSE-FACTOR1. Gene Dev 12: 3703–3714.

79. Sugano, S.S., Shimada, T., Imai, Y., Okawa, K., Tamai, A., Mori, M., and Hara-Nishimura, I. (2010). Stomagen positively regulates stomatal density in Arabidopsis. Nature 463: 241–244.

80. Sun, J., Nishiyama, T., Shimizu, K., and Kadota, K. (2013). TCC: an R package for comparing tag count data with robust normalization strategies. Bmc Bioinformatics 14: 219.

81. Takeda, K. and Shibaoka, H. (1981). Changes in microfibril arrangement on the inner surface of the epidermal cell walls in the epicotyl of Vigna angularis ohwi et ohashi during cell growth. Planta 151: 385–392.

82. Tanaka, Y., Sugano, S.S., Shimada, T., and HaraDNishimura, I. (2013). Enhancement of leaf photosynthetic capacity through increased stomatal density in Arabidopsis. New Phytol 198: 757–764.

83. Tang, Y., Horikoshi, M., and Li, W. (2016). ggfortify: Unified Interface to Visualize Statistical Results of Popular R Packages. R Journal 8: 474–485.

84. Thomas, S.G., Phillips, A.L., and Hedden, P. (1999). Molecular cloning and functional expression of gibberellin 2-oxidases, multifunctional enzymes involved in gibberellin deactivation. Proc National Acad Sci USA 96: 4698–4703.

85. Tsuge, T., Tsukaya, H., and Uchimiya, H. (1996). Two independent and polarized processes of cell elongation regulate leaf blade expansion in *Arabidopsis thaliana* (L.) Heynh. Development 122: 1589–600.

86. Tsukaya, H. (2018). Leaf shape diversity with an emphasis on leaf contour variation, developmental background, and adaptation. Semin Cell Dev Biol 79: 48–57.

87. Veen, H., Akman, M., Jamar, D.C.L., Vreugdenhil, D., Kooiker, M., Tienderen, P., Voesenek, L.A.C.J., Schranz, M.E., And Sasidharan, R. (2014). Group VII Ethylene Response Factor diversification and regulation in four species from flood prone environments. Plant Cell Environ 37: 2421–2432.

88. Voesenek, L.A.C.J. and Sasidharan, R. (2013). Ethylene – and oxygen signalling – drive plant survival during flooding. Plant Biology 15: 426–435.

89. Wanke, D. (2011). The ABA-mediated switch between submersed and emersed life-styles in aquatic macrophytes. J Plant Res 124: 467–475.

90. Wells, C.L. and Pigliucci, M. (2000). Adaptive phenotypic plasticity: the case of heterophylly in aquatic plants. Perspect Plant Ecol Syst 3: 1–18.

91. Wickham, H. et al. (2019). Welcome to the Tidyverse. J Open Source Softw 4: 1686.

92. Xu, K., Xu, X., Fukao, T., Canlas, P., Maghirang-Rodriguez, R., Heuer, S., Ismail, A.M., Bailey-Serres, J., Ronald, P.C., and Mackill, D.J. (2006). Sub1A is an ethylene-response-factor-like gene that confers submergence tolerance to rice. Nature 442: 705–708.

93. Xu, P., Chen, H., Ying, L., and Cai, W. (2016). AtDOF5.4/OBP4, a DOF Transcription Factor Gene that Negatively Regulates Cell Cycle Progression and Cell Expansion in *Arabidopsis thaliana*. Sci Rep 6: 27705.

94. Zhang, L.-Y. et al. (2009). Antagonistic HLH/bHLH Transcription Factors Mediate Brassinosteroid Regulation of Cell Elongation and Plant Development in Rice and Arabidopsis. Plant Cell 21: 3767–3780.

