## Supplemental Information for "The molecular framework of heterophylly in *Callitriche palustris* L. differs from that in other amphibious plants"

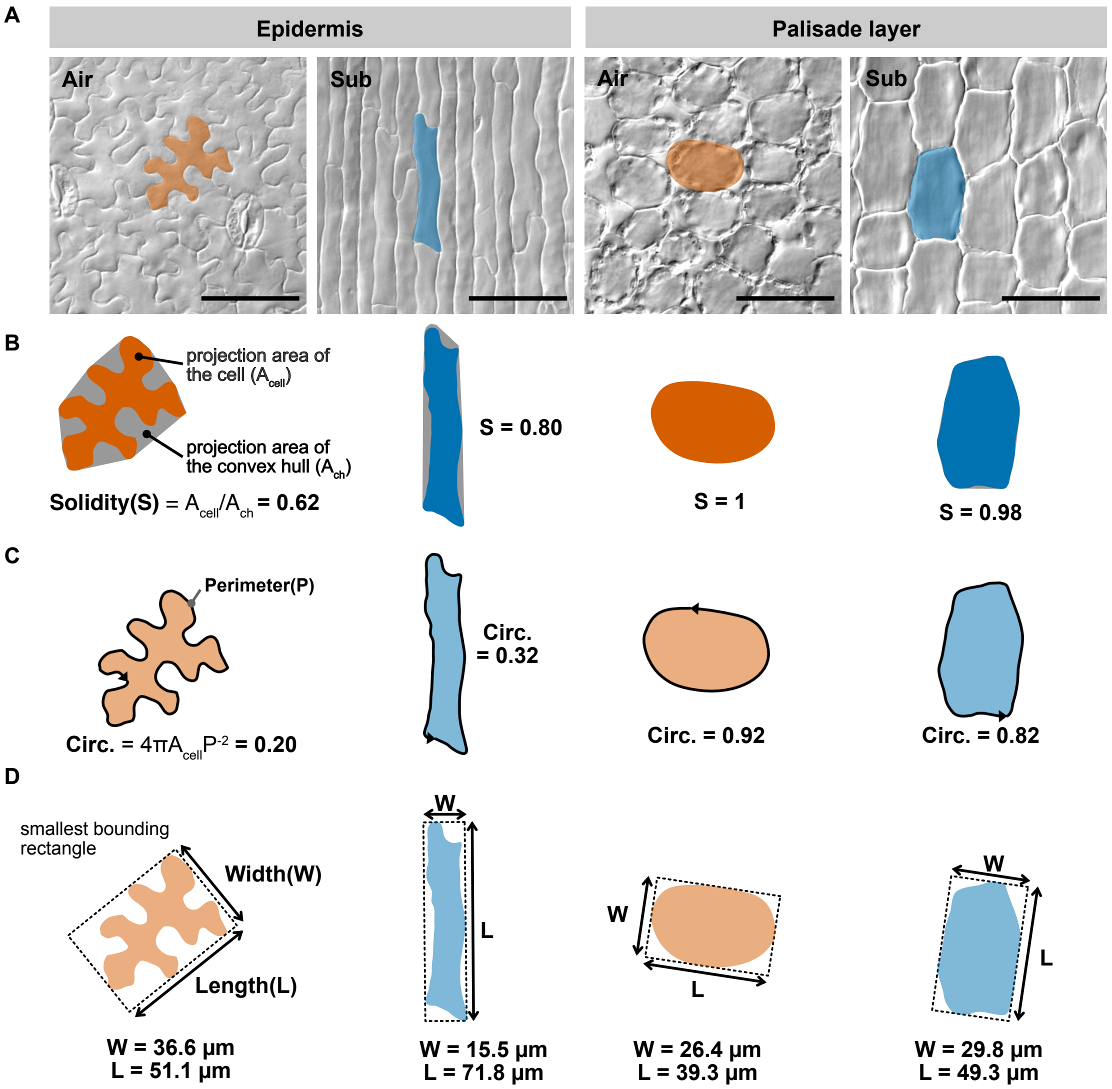


**Supplemental Figure 1. Explanation scheme of cell morphology measurement.** (Supports Figures 1, 2, and 3)

Some cell shape indices used in this study are schematized using actual cell data. (A) The projected cell shape was traced from the original images. (B-D) Measurement schemes of (B) solidity, (C) circularity, and (D) cell width and length. Considering the observed shape of the cells of aerial and submerged leaves, we analyze the cell shape with multiple indices rather than with just one index. For example, circularity, which (or its reciprocal) is often used for indicator of the complexity of the contour, can be reduced not only by the higher complexity of the contour, but also by the elongated shape. Scale bars: 50 μm.

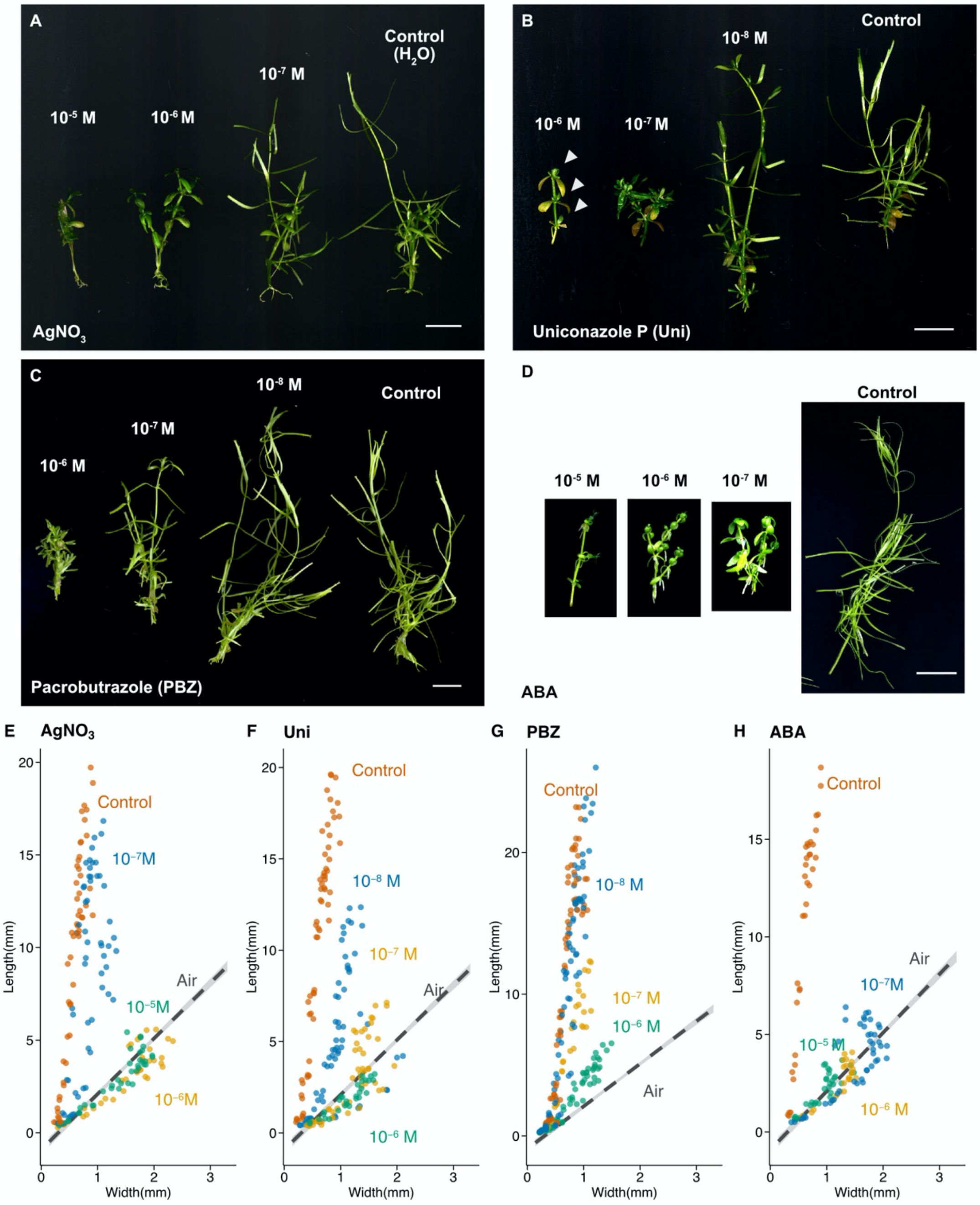


**Supplemental Figure 2. Effects of phytohormone inhibitors on *C. palustris* grown under submerged conditions**. (Supports Figure 2)

(A–D) Images of whole plants treated with various concentrations of the phytohormone inhibitors (A) $AgNO_3$, (B) uniconazole P, and (C) paclobutrazol or the phytohormone (D) ABA under submerged conditions. Arrowheads indicate substantially inhibited shoot growth. (E–H) Length–width plots of leaves collected from 3–4 shoots, from 2–3 biological replicates. Plots of leaves treated with various concentrations of (E) $AgNO_3$, (F) uniconazole P, (G) paclobutrazol, or (H) ABA. Gray lines represent the regression lines of normal aerial shoots from a recent study (Koga et al., 2020) and are included for comparison purposes. Distilled water was used as the control treatment for (A), (D), (E), and (H), whereas 0.1% ethanol was used as the control treatment for (B), (C), (F), and (G). Scale bars: 1 cm.

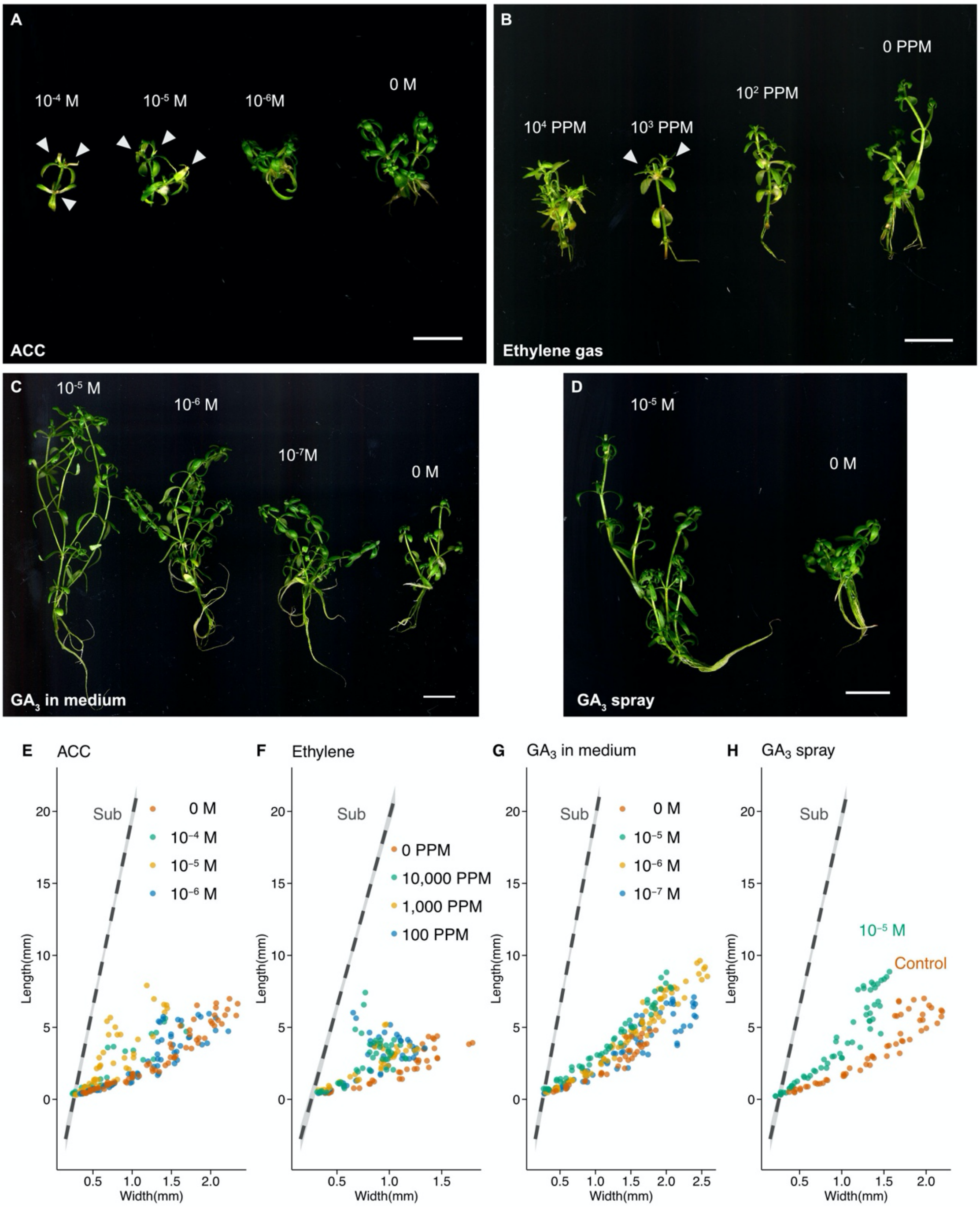


**Supplemental Figure 3. Effects of phytohormones on *C. palustris* under aerial growth conditions.** (Supports Figure 3)

(A–D) Images of whole plants treated with various concentrations of phytohormones: (A) ACC supplied in the growth medium, (B) ethylene gas, (C) $GA_3$ supplied in the growth medium, and (D) $GA_3$ sprayed under aerial conditions. (E–H) Length–width plots of leaves collected from 3–4 shoots from 2–3 biological replicates. Plots of leaves treated with various concentrations of (E) ACC, (F) ethylene gas, (G) $GA_3$, and (H) $GA_3$ (spray application). Gray lines represent the regression lines of normal submerged shoots from a recent study (Koga et al., 2020) and are included for comparison purposes. Scale bars: 1 cm.

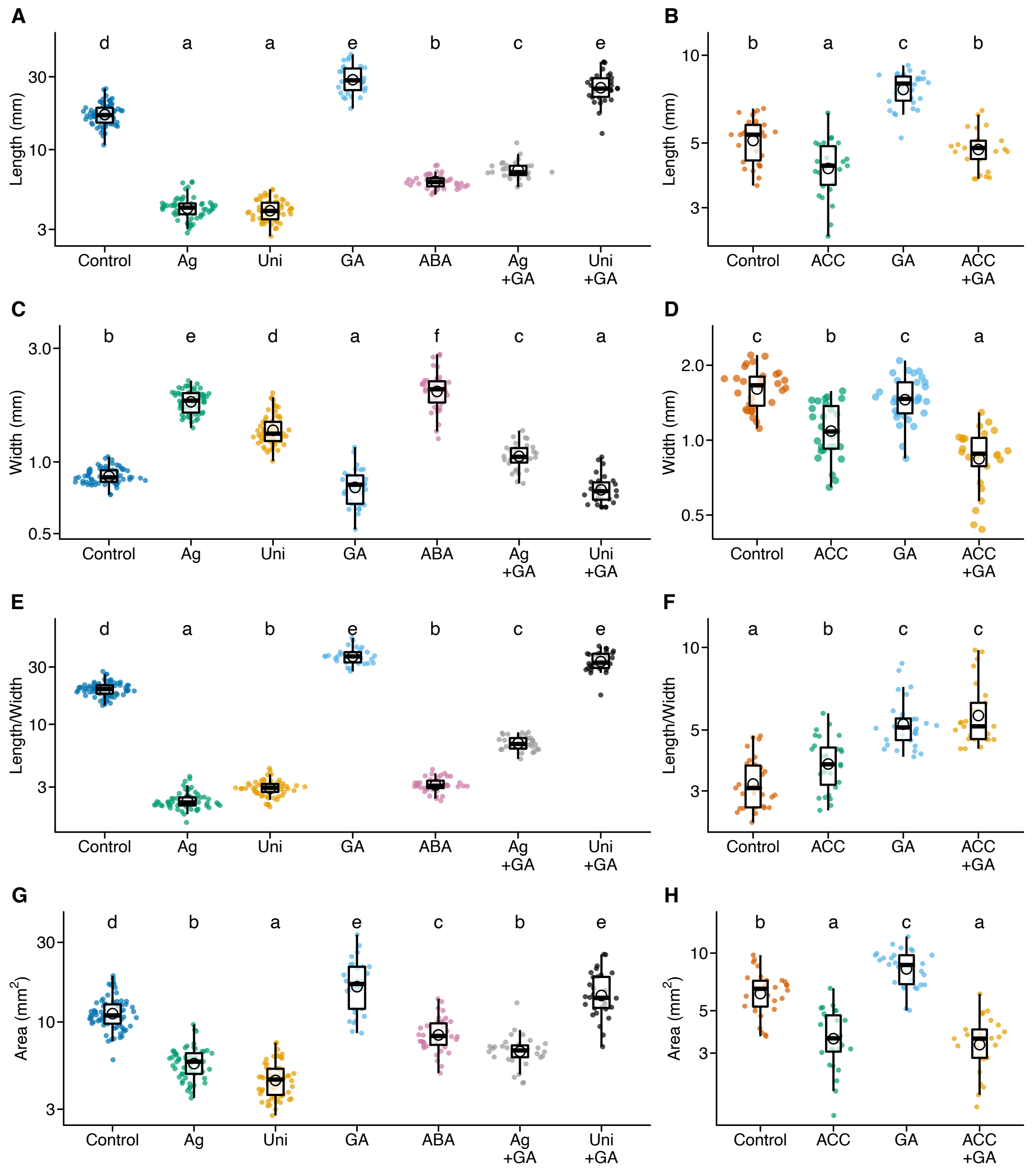


**Supplemental Figure 4**. ***Callitriche palustris*** **leaf shapes after hormone treatments.** (Supports Figures 2 and 3)

Plots of (A, B) leaf length, (C, D) leaf width, (E, F) leaf index (length/width), and (G, H) leaf area of mature leaves from plants treated with hormones and inhibitors. (A, C, E, G) Treatments under submerged conditions and (B, D, F, H) treatments under aerial conditions. Significant differences ($P < 0.05$, Tukey's test) are indicated with different letters.

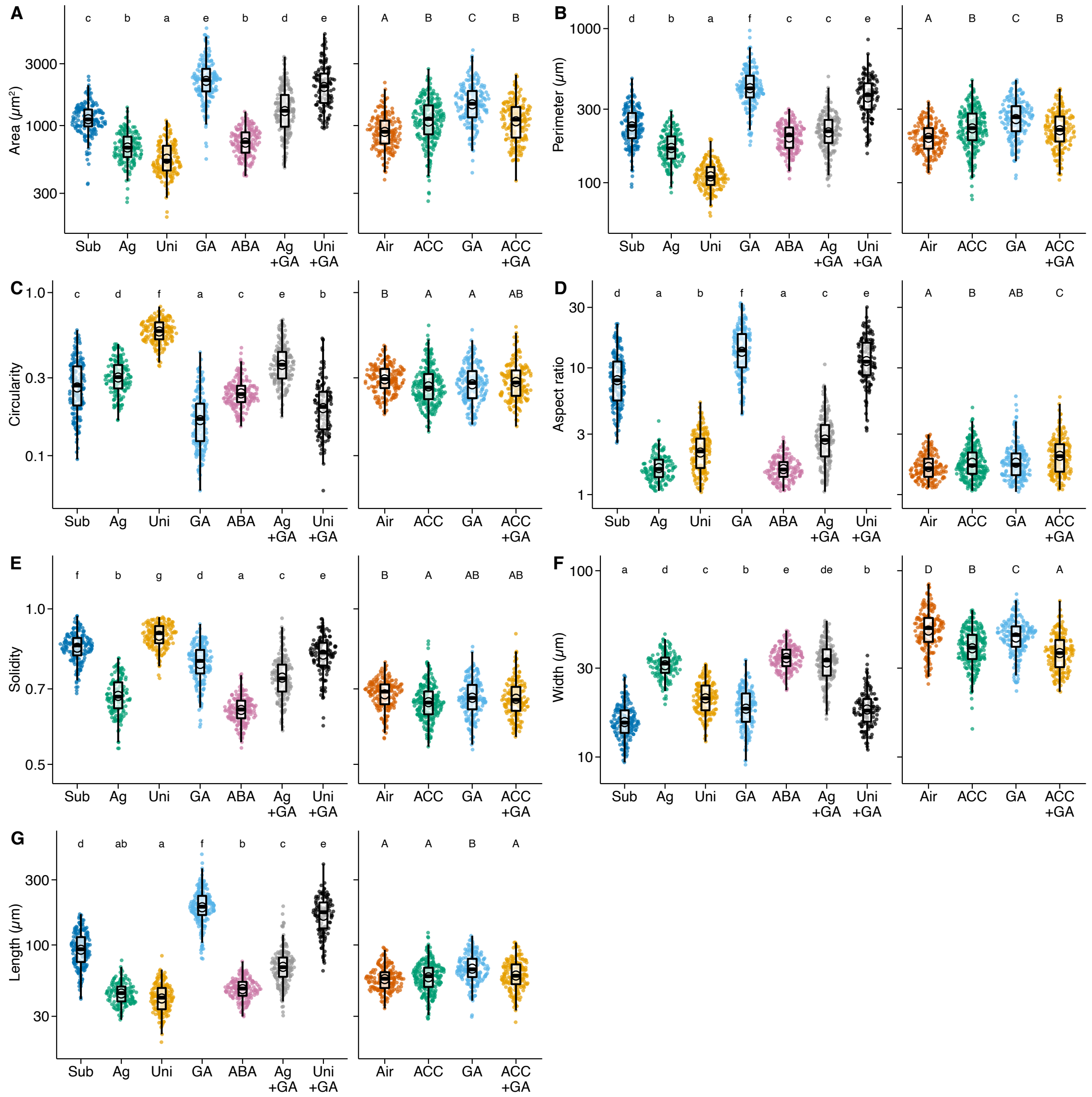


**Supplemental Figure 5. Changes in pavement cells following hormone/inhibitor treatments of *C. palustris.*** (Supports Figures 2 and 3)

Plots for cell area (A), perimeter (B), circularity (C), aspect ratio of the fitted ellipse (D), solidity (E), cell width (F), and cell length (G). Significant differences ($P < 0.05$, Tukey's test) are indicated with different letters.

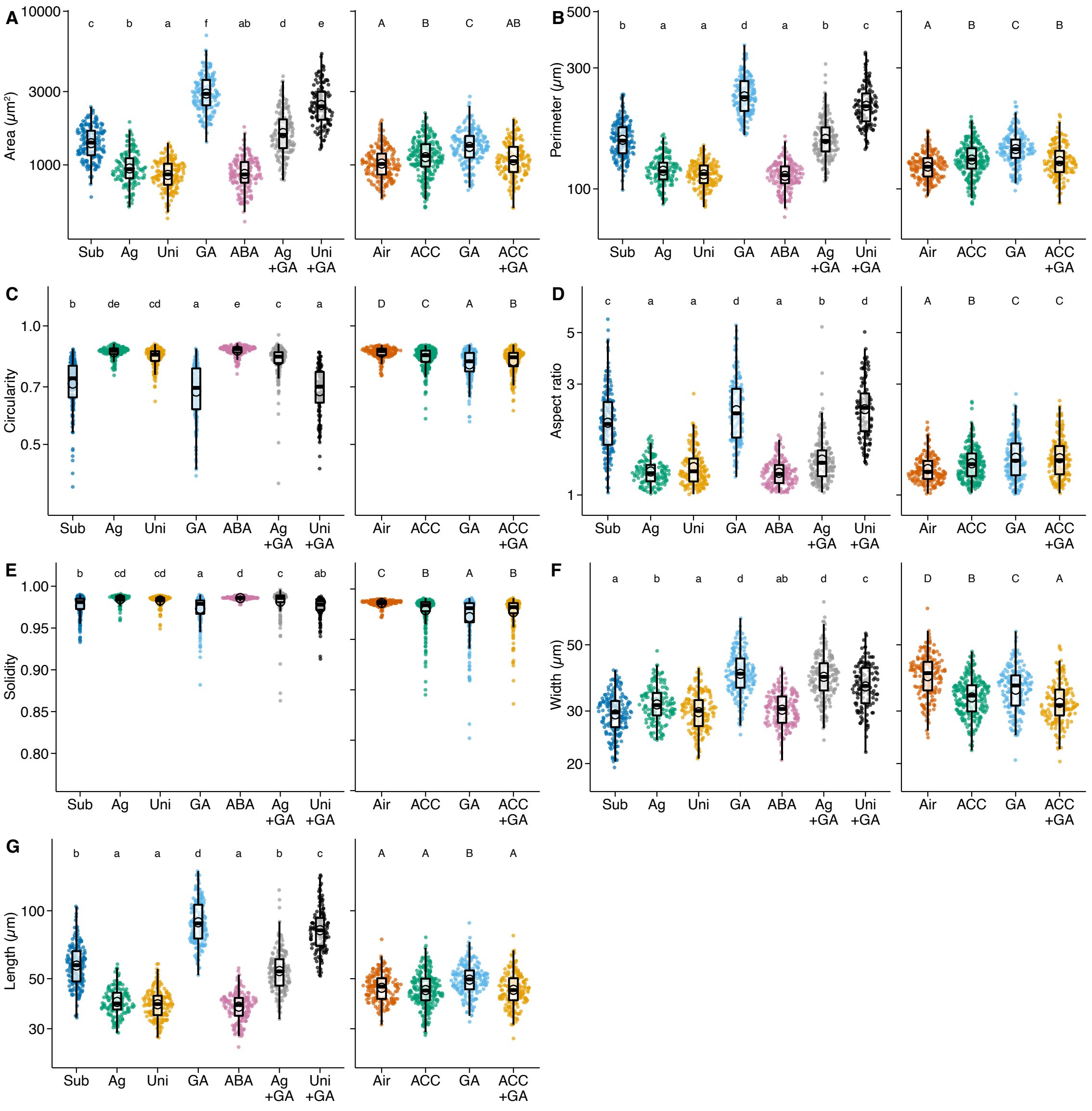


**Supplemental Figure 6. Changes in palisade cells following hormone/inhibitor treatments of *C. palustris*.** (Supports Figures 2 and 3)

Plots for cell area (A), perimeter (B), circularity (C), aspect ratio of the fitted ellipse (D), solidity (E), cell width (F), and cell length (G). Significant differences ($P < 0.05$, Tukey's test) are indicated with different letters.

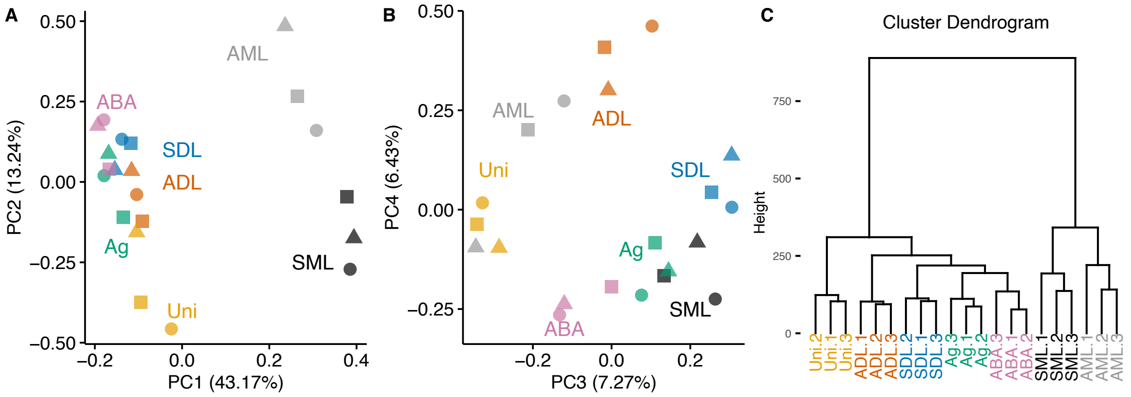


**Supplemental Figure 7. Comparison of transcriptome profiles in *C. palustris* samples.**

(Supports Figure 4)

(A, B) Principal component analysis of RNA-seq data. Different shapes denote individual plants. (C) A cluster dendrogram of hierarchical clustering using Euclidean distances and Ward's method. $Log_2$-transformed TPM values of 66,905 expressed genes were used. Colors indicate different samples. ADL: aerial leaf primordia, SDL: submerged leaf primordia, Ag: $AgNO_3$-treated submerged leaf primordia, Uni: uniconazole P-treated submerged leaf primordia, ABA: ABA-treated submerged leaf primordia, AML: aerial mature leaves, SML: submerged mature leaves.

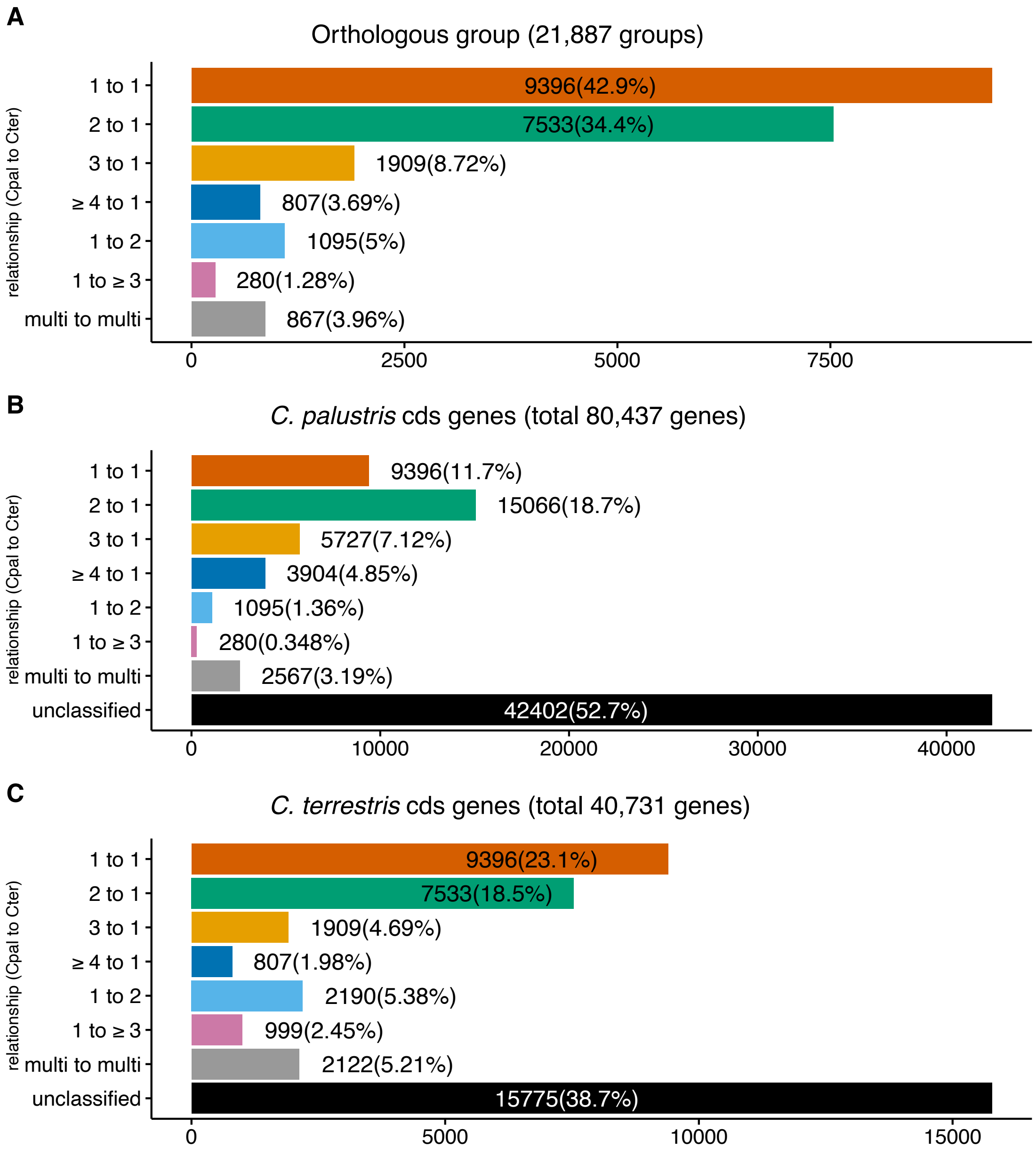


**Supplemental Figure 8. Orthologous relationship distribution in *C. palustris* and *C. terrestris* coding genes.** (Supports Figures 5)

*C. palustris* and *C. terrestris* coding genes were assigned to ortholog groups by orthofinder, then each orthologous group was classified to one of the orthologous relationships based on the number of genes in a group. For example, “2 to 1” relationship represent that *C. palustris* has two orthologs while *C.terrestris* has only one counterpart.

(A) Distribution of relationship in the orthologous groups. (B, C) Numbers of (B) *C. palustris* and (C) *C. terrestris* genes in the ortholog groups. “unclassified” represents genes in which orthology was not determined by orthofinder, either because they have no counterpart in the other species, or because of erroneous assembly or cds misprediction. Nearly half (46.8%) ortholog groups include two or more paralogs in *C. palustris,* but only one ortholog in *C.terrestis.*

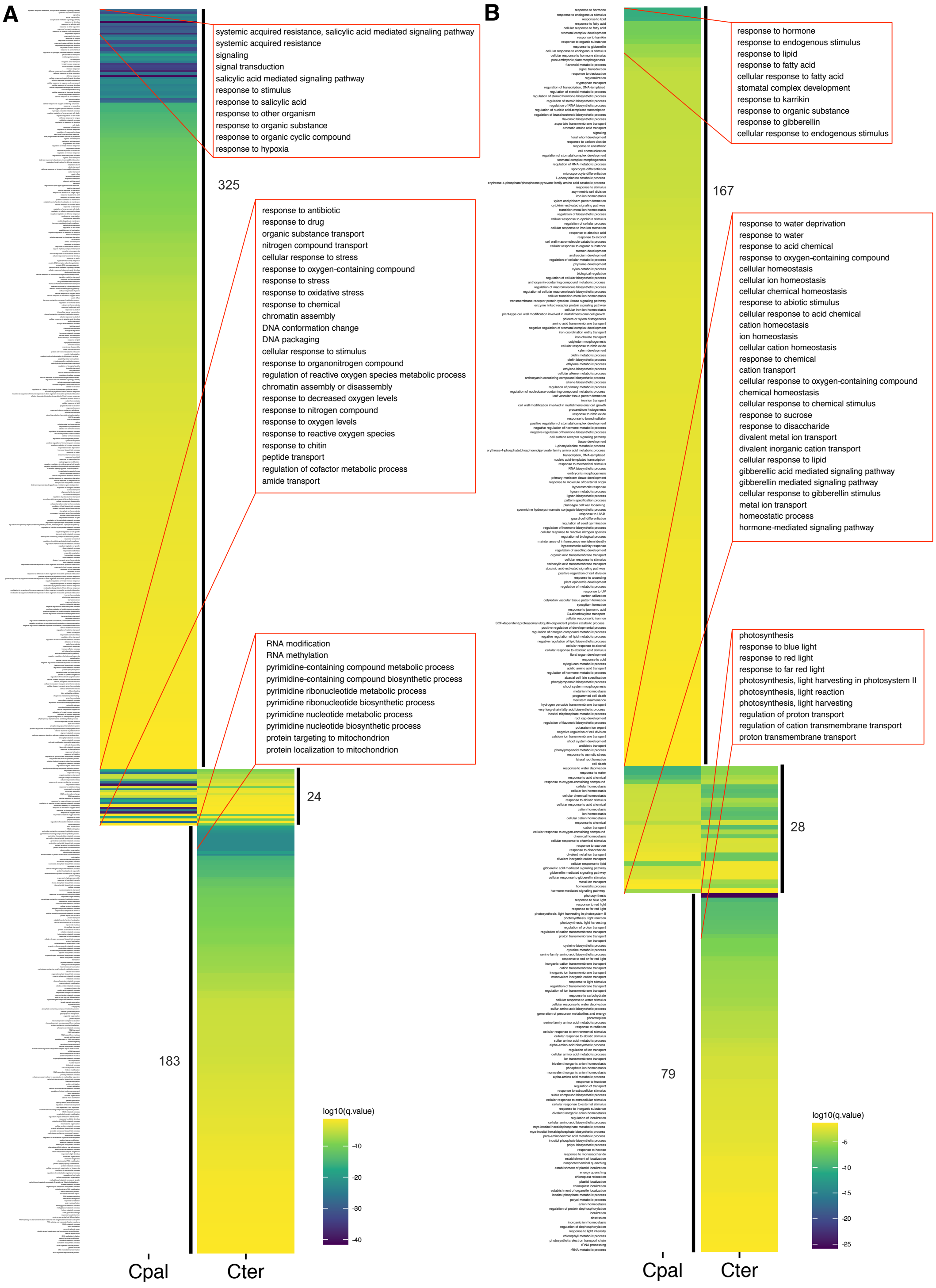


**Supplemental Figure 9. Comparison of significantly enriched GO terms between the DEGs of *C. palustris* and *C. terrestris* leaf primordia.** (Supports Figures 5)

Significantly enriched (adjusted $P < 0.05$) biological process GO terms among the DEGs between the aerial and submerged leaf primordia of *C. palustris* and *C. terrestris*. (A) Enriched terms among the upregulated DEGs in submerged leaf primordia. (B) Enriched terms among the downregulated DEGs in submerged leaf primordia. Ten highly significant but non-overlapping terms as well as all overlapping terms between *C. palustris* and *C. terrestris* are listed in boxes.

**Supplemental Table 1.** ***de novo*** **transcriptome assembly status**

| **Species** | | *C. palustris* | *C. terrestris* |
|---|---|---|---|
| **# of genes** | | 145,929 | 74,145 |
| **# of contigs** | | 243,784 | 130,314 |
| **Total base** | | 99,628,636 | 62,970,934 |
| **N50 (genes)** | | 1,040 bp | 1,229 bp |
| **Mean (genes)** | | 721 bp | 757 bp |
| **BUSCO v4 Eudicots** | **Complete (Single-copy)** | 74.7% | 86.4% |
| | **Complete (Duplicated)** | 13.6% | 1.9% |
| | **Fragment** | 4.5% | 3.8% |
| | **Missing** | 7.0% | 7.9% |
| **# of coding genes** | | 80,437 | 40,731 |
| **# of orthologs decided** | | 34,083 | 19,644 |

**Supplemental Data Set 1. Sequence and mapping results**

(supplied as a separate file)

**Supplemental Data Set 2. Significantly enriched GO term lists**

(supplied as a separate file)

**Supplemental Data Set 3. Measurement of hormonal contents**

(supplied as a separate file)

**Supplemental Data Set 4. Annotations of the candidate gene set**

(supplied as a separate file)

**Supplemental Data Set 5. Enriched GO term lists in the candidate gene set**

(supplied as a separate file)
